# Species-dependent clearance of alternariol monomethyl ether, tenuazonic acid and altertoxin II in rat and human primary hepatocytes

**DOI:** 10.64898/2026.07.13.737671

**Authors:** Yuri Bastos-Moreira, Charlène Gendre, Florian Call, Daniel Piller, Eva Waltenberger, Andreas Wasilewicz, Judith M. Rollinger, Jérôme Henri, Doris Marko, Ludovic Le Hégarat, Elisabeth Varga

**Affiliations:** Department of Food Chemistry and Toxicology, Faculty of Chemistry, University of Vienna, Währinger Str. 38, 1090 Vienna, Austria; ANSES, French Agency for Food, Environmental and Occupational Health & Safety, Fougères Laboratory, Toxicology of contaminants unit, 10 B rue Claude Bourgelat, 35306 Fougères, France; Doctoral School in Chemistry, Faculty of Chemistry, University of Vienna, Währinger Str. 42, 1090 Vienna, Austria; Division of Pharmacognosy, Department of Pharmaceutical Sciences, University of Vienna, Josef-Holaubek-Platz 2, Vienna 1090, Austria; Vienna Doctoral School of Pharmaceutical, Nutritional and Sport Sciences, University of Vienna, Vienna, Austria; ANSES, French Agency for Food, Environmental and Occupational Health & Safety, Fougères Laboratory, Kinetics, Metabolism Modelling and Experiments Unit, 10 B rue Claude Bourgelat, 35306 Fougères, France; Food Hygiene and Technology, Centre for Food Science, Clinical Department for Farm Animals and Food System Transformation, University of Veterinary Medicine, Vienna, Veterinärplatz 1, 1210 Vienna, Austria

**Keywords:** *Alternaria alternata*, toxicokinetics, intrinsic clearance, hepatic metabolism, LC-MS/MS

## Abstract

The occurring food contaminants alternariol monomethyl ether (AME), tenuazonic acid (TeA) and altertoxin II (ATX-II) are recognized as emerging *Alternaria* mycotoxins, yet data gaps remain regarding their toxicokinetic characteristics. The hepatic metabolism of these three substances was investigated in primary rat (PRH) and human (PHH) hepatocytes by monitoring parent compound depletion and, where applicable, metabolite formation. AME was initially evaluated at 1 µM over 4 h and subsequently investigated in concentration-dependent experiments (0.75, 1.5, 3, and 8 µM). TeA was assessed at 5 µM over 4 h, whereas ATX-II was evaluated at 1.11 µM over 4 h and at 0.22 µM over 30 min. For AME, time-dependent depletion was further investigated in PRHs at two hepatocyte densities (0.25 and 0.5 × 10^6^ cells/mL). In PRHs, AME metabolism followed Michaelis–Menten kinetics (V_max_ = 150.9 pmol·min^-¹^·10^-6^ cells, K_m_ = 1.18 µM), whereas no reliable kinetic model could be established for PHHs. In contrast, TeA exhibited high metabolic stability, with only 9–10% depletion after 4 h, indicating negligible hepatic clearance in both species. ATX-II was also rapidly depleted and became undetectable within 30 min, accompanied by formation of altertoxin I (ATX-I), which was more pronounced in PHHs than in PRHs. Substrate depletion revealed interspecies differences in hepatic clearance capacity and stability. Furthermore, tentative metabolite analysis indicated that AME undergoes phase I and phase II biotransformation. Overall, these findings provide comparative insights into PRH and PHH systems, offering a foundation for future studies on their toxicological relevance and impact on human health.

## 1. Introduction

Fungal contamination of food and feed represents a major challenge, threatening food safety and agricultural productivity globally. Among the fungal genera most frequently associated with the contamination and spoilage of agricultural commodities are *Aspergillus*, *Penicillium*, *Fusarium*, and *Alternaria* (Crudo et al., 2019). Species of the genus *Alternaria* are ubiquitous phytopathogens and saprophytes that infect a wide range of crops and contaminate cereals, fruits, vegetables, and their derived food products both before and after harvest (Aichinger et al., 2021). In addition to causing substantial crop losses and associated economic impacts, *Alternaria* species produce a diverse array of secondary metabolites. More than 70 of these compounds have been reported to exert adverse effects in vertebrates and are therefore collectively referred to as *Alternaria* toxins. These compounds have been extensively studied and shown to exert a broad spectrum of adverse biological effects in both *in vitro* and *in vivo* models, including acute toxicity, genotoxicity, mutagenicity, carcinogenicity, estrogenic activity, and immunomodulatory effects (Crudo et al., 2019; Louro et al., 2024).

Several mycotoxins, including aflatoxins produced by *Aspergillus* spp. and fumonisins produced by *Fusarium* spp., are regulated under Commission Regulation (EU) 2023/915 and amendments (European Commission, 2023). In contrast, *Alternaria* mycotoxins are currently classified as “emerging mycotoxins” owing to the limited availability of toxicological and occurrence data and are therefore not subject to legally binding maximum limits. Nevertheless, in recognition of their potential risk to public health, the European Commission issued a recommendation in April 2022, encouraging the monitoring of the most prevalent *Alternaria* toxins in food and the investigation of factors contributing to elevated contamination levels (European Medicines Agency, 2023).

The chemical structures of selected *Alternaria* mycotoxins, grouped according to their structural characteristics, are shown in **Fig. 1**. Previous studies have demonstrated that these toxins exhibit genotoxic activity through distinct mechanisms of action. Alternariol monomethyl ether (AME), a dibenzo-α-pyrone derivative, induces DNA strand breaks through inhibition of human topoisomerases I and II and the generation of oxidative stress (Fehr et al., 2009; Solhaug et al., 2016). In contrast, the epoxide-containing perylene quinone altertoxin II (ATX-II) is believed to exert its genotoxic effects primarily through the formation of covalent DNA adducts (Soukup et al., 2020). Tenuazonic acid (TeA), an amine/amide metabolite, is the most frequently reported *Alternaria* toxin in food samples (Marin et al., 2013). Although TeA exhibits relatively low acute toxicity and no confirmed evidence of chronic toxicity, it remains toxicologically relevant because of its frequent occurrence in foods (EFSA et al., 2016).

**Figure 1:**
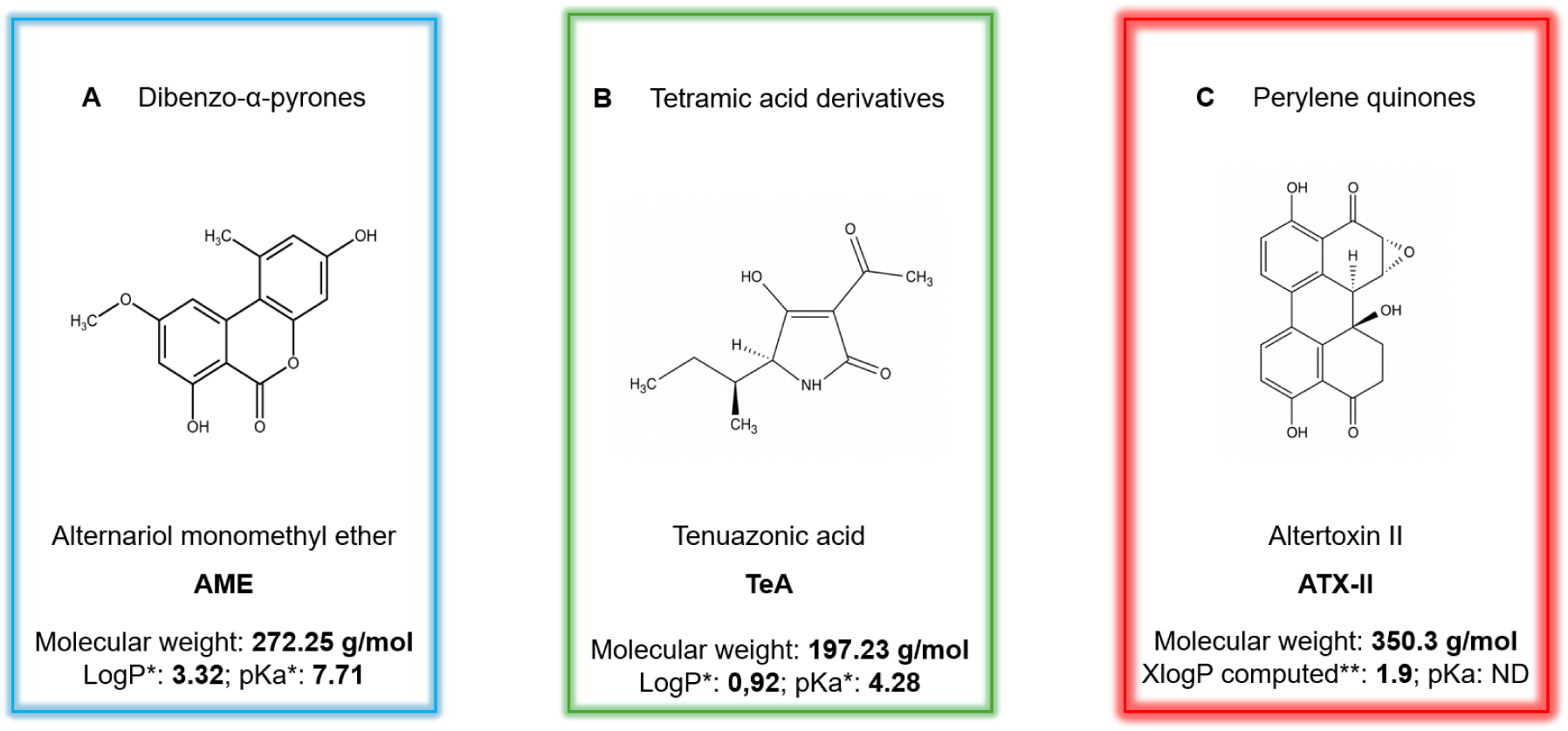
Chemical structures of selected *Alternaria* mycotoxins grouped according to their structural classes: A) dibenzo-α-pyrones like alternariol monomethyl ether (AME); B) tetramic acid derivatives like tenuazonic acid (TeA); C) perylene quinones like altertoxin II (ATX-II). LogP and pKa values are provided where available. *LogP and pKa values were obtained from Tölgyesi *et al*. (2015). **XlogP values were calculated using XLogP3 3.0 (PubChem, release 2025.04.14). ND, not determined.

**Figure 1.**
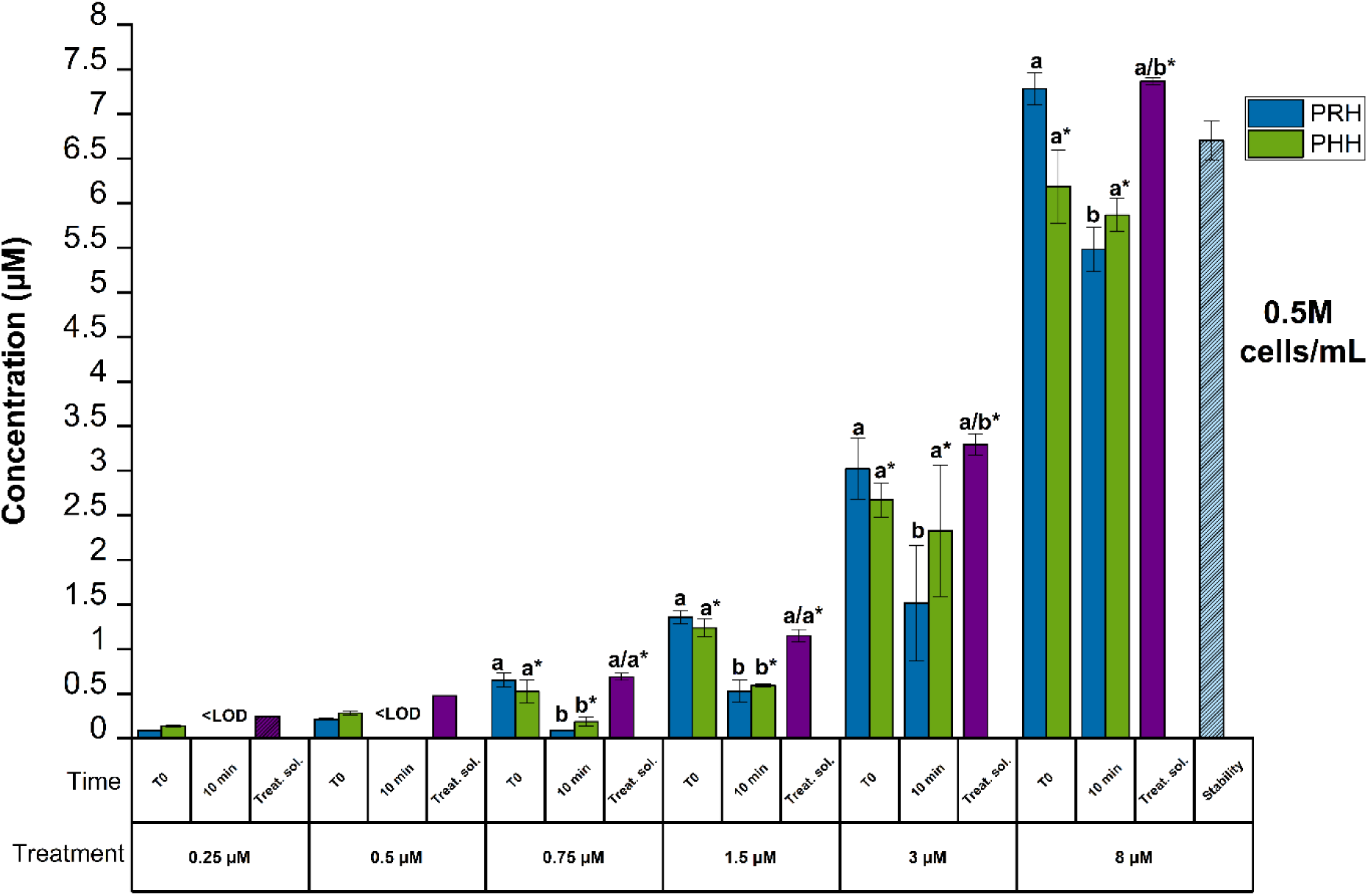
The concentration of alternariol monomethyl ether (AME) in primary rat (PRH) and human (PHH) hepatocytes following treatment at increasing concentrations (0.25–8 µM). Stability controls were prepared in the same manner as the kinetic samples by adding 100 µL of treatment solution to 100 µL of DMSO-free medium (1:1, *v/v*) and incubating at 37 °C with 5 % carbon dioxide (CO_₂_) for the full duration of the experiment in the absence of hepatocytes. Treat. sol. denotes treatment solution controls prepared by mixing 100 µL of treatment solution with 100 µL of DMSO-free medium (1:1, *v/v*), followed by immediate extraction without incubation. The AME concentrations were quantified using liquid chromatography tandem mass spectrometry (LC-MS/MS). Results are depicted as mean ± standard deviation (SD) of three technical replicates. Statistically significant differences in AME concentrations among multiple groups (≥3) were evaluated by applying one-way ANOVA, followed by Fisher’s LSD as a post hoc test. Comparison between two groups was performed using the Student’s *t*-test. Different letters indicate significant differences (*p* < 0.05) between T0 and 10 min. * Corresponds to PHHs.

In addition to their toxicological properties, several *Alternaria* mycotoxins exhibit considerable chemical stability. Significant proportions of these compounds remain intact following thermal food processing and simulated gastrointestinal digestion (Call et al., 2026). Their frequent co-occurrence in food matrices, together with the possibility of additive or synergistic toxicological interactions, highlights the need for comprehensive occurrence data and improved approaches to risk assessment.

The present study aimed to investigate the hepatic clearance and metabolic stability of AME, TeA, and ATX-II in primary rat hepatocytes (PRHs) and primary human hepatocytes (PHHs) using targeted liquid chromatography–tandem mass spectrometry (LC-MS/MS). Specifically, time- and concentration-dependent clearance of AME, the metabolic stability of TeA, and the depletion of ATX-II together with the formation of its metabolite altertoxin I (ATX-I) were evaluated. The findings provide new insights into the inter-species clearance of major *Alternaria* mycotoxins and strengthen the toxicological database needed to support future human health risk assessment.

## 2. Materials and methods

### 2.1. Chemicals and reagents

#### 2.1.1 Toxicokinetics

AME was synthesized according to Mikula *et al*. (2013) and kindly provided by Prof. Roderich Süssmuth (Technical University of Berlin, Berlin, Germany). Tenuazonic acid (TeA) was purchased from SantaCruz Biotechnology (SC-202357A, Dallas, Texas, USA) and the altertoxin II (ATX-II) was isolated from an *Alternaria* culture extract and purified ourselves, detailed information on this reference material can be found in **section 2.1.1.1**. Dimethyl sulfoxide (DMSO) was purchased from Sigma (St.-Quentin-Fallavier, France). The AME stock solution was prepared at a concentration of 10 mM in DMSO, the TeA stock solution was prepared at 20 mM in DMSO and the ATX-II stock solution was prepared at 10 mM in ethanol, all stock solutions were stored at –20 °C. To stop metabolism, acetonitrile (ACN) and methanol (MeOH) (LC-MS grade) were purchased from Thermo Fisher Scientific (Waltham, MA, USA)

#### 2.1.1.1 Alternaria extract production and ATX-II isolation

Sterile mCDB medium (4% glucose, 0.1% KH_2_PO_4_, 0.1% NaNO_3_, 0.1% yeast extract, 0.05% MgSO_4_ 7 x H_2_O, 0.025% KCl, 0.025% NaCl, 0.025% NH_4_Cl, 0.001% FeSO_4_ 7 x H_2_O, 0.001% ZnSO_4_) was inoculated with mycelium of *Alternaria alternata* strain ATCC66981 and cultivated for 7 days at 28 °C in the dark. Following cultivation, the fungal culture was extracted with ethyl acetate at a ratio of 1:2 (culture: ethyl acetate, *v/v*) under continuous shaking for 1 h. The organic phase was subsequently collected, and the solvent was removed under reduced pressure at 4 °C using a SpeedVac concentrator (Laconco).

ATX-II was then isolated from the *Alternaria* culture extract by reversed-phase flash chromatography using a puriFlash 4250 system (Interchim, Montluçon, France) equipped with Evaporative Light Scattering Detector, Photodiode Array detection and a fraction collector. The extract (200 mg) was fractionated on a C18-HQ column using a water/acetonitrile gradient (5–98% acetonitrile), and ATX-II-containing fractions were identified by Thin-Layer Chromatography, pooled, and yielded 6.22 mg of purified ATX-II. Compound identity was confirmed by 1H and 13C Nuclear Magnetic Resonance (NMR) and 2D NMR (COSY, HSQC, and HMBC), with spectroscopic data consistent with previously reported data for ATX-II (Mahmoud et al., 2022). More information on the isolation, purification, and structural confirmation of ATX-II from that extract can be found in **Supplementary Information I**.

#### 2.1.2 Primary hepatocyte source and required media with supplements

Cryopreserved (Sprague-Dawley) pool PRHs (Grade S for suspension assays, batch HEP134070) (six animals were used to generate the hepatocyte pool from male rats) and cryopreserved pool PHHs (10 Caucasian donor mixed gender pooled: 5 men and 5 women) (Grade S for suspension assays, batch HEP190021-TA05) were purchased from Wepredic (Biopredic International, Saint-Grégoire, France). Cellular metabolic functions were validated by the supplier and measured by LC-MS/MS for the following phase I enzyme reference substrates: CYP1A2, CYP2B6, CYP2D6, and CYP3A4/5, as well as for the following phase II-dependent activities: UGT1A1, UGT1A6, UGT1A7, and SULT1A. Thawing medium (MIL130), treatment medium is composed of basal hepatic cell medium (MIL600) supplemented with an aliquot of additives (ADD222) composed of penicillin 100 UI/mL, streptomycin 100 µg/mL, bovine insulin 4 µg/mL and hydrocortisol 50 µM (ADD222) were purchased from Wepredic (Biopredic International, Saint-Grégoire, France).

#### 2.1.3. LC-MS/MS

ACN, MeOH, and LC-MS/MS water were purchased from Honeywell (Seelze, Germany). Ammonium acetate and ammonium hydroxide were obtained from Sigma-Aldrich (Vienna, Austria), while DMSO was purchased from Carl Roth GmbH + Co. KG (Karlsruhe, Germany). For targeted LC-MS/MS, reference standards for AME, TeA and ATX-I were purchased from Romer Labs Diagnostic GmbH (Tulln, Austria). As ATX-I is the primary reductive metabolite of ATX-II, it was included in the targeted LC–MS/MS analysis to monitor the biotransformation of ATX-II. Since an analytical standard for ATX-I was available, its formation could be quantified alongside the depletion of ATX-II.

### 2.2 Treatment of primary hepatocytes with AME, TeA and ATX-II

#### 2.2.1 Kinetics with PRHs and PHHs

The cryopreserved vials of pool hepatocytes were thawed in a water bath at 37 °C and transferred to a thawing medium heated to 37 °C (MIL130), according to supplier recommendations. After centrifugation at 100 g and 180 g for rat and human hepatocytes, respectively, the supernatant was removed, and the cells were re-suspended in 3 mL treatment medium. Viability after reconstitution was determined using Trypan blue at 0.4% and was always above 80%. Then, PRHs and PHHs were incubated for independent kinetics in suspension in 5 mL glass tubes (to minimize nonspecific binding to plastic surfaces), at a final density of 5.10^5^ cells/mL in a total volume of 200 µL containing 0.05% DMSO for AME and TeA or ethanol for ATX-II (final concentration). Samples were maintained at 37 °C in a humidified atmosphere of 5% CO_2_ with continuous agitation on a microplate shaker. At the end of the incubation period, cell viability was assessed by using the CellTiter-Glo^®^ Luminescent Cell Viability Assay (Promega Corporation, Madison, WI, USA), that measures ATP content.

#### 2.2.1.1 AME kinetics

Primary rat and human hepatocytes in suspension were first incubated at final density of 5.10^5^ cells/mL with AME at a non-toxic concentration of 1 µM in 0.05% of DMSO in triplicate at the following time points: time zero (T0), 30 min, 1, 1.5, 2, and 4 h, independently. Next, a second experiment was conducted using only PRHs in the presence of AME at non-cytotoxic concentration of 5 µM in 0.05% of DMSO at time points T0, 5, 10, 15, and 30 min, at two different cell densities: 0.25 and 0.5 million cells per milliliter, independently, in order to optimize cell density for accurate assessment of AME depletion. A third and final experiment yielded the greatest disappearance of AME; Michaelis–Menten parameters (V_max_ and K_m_) were calculated using PRHs and PHHs, with AME at 6 different non-cytotoxic concentrations in 0.05% of DMSO (0.25, 0.5, 0.75, 1.5, 3, and 8 µM) for time points of 0 and 10 min. For each time point, three technical replicates were performed.

#### 2.2.1.2 TeA kinetics

Primary rat and human hepatocytes in suspension at final density of 5.10^5^ cells/mL were incubated with TeA at a non-toxic concentration of 5 µM in 0.05% of DMSO for T0, 30 min, 1, 1.5, 2, and 4 h, independently. Three technical replicates were performed.

#### 2.2.1.3 ATX-II kinetics

Primary rat and human hepatocytes in suspension at final density of 5.10^5^ cells/mL were first incubated with ATX-II at a non-cytotoxic concentration of 1.11 µM in 0.05% of ethanol for the following durations: T0, 30 min, 1, 1.5, 2, and 4 h, independently. A second experiment was performed using primary rat and human hepatocytes with ATX-II at 0.22 µM in 0.05% of ethanol at the following time points: T0, 5, 10, 20 and 30 min. For each time point, three technical replicates were performed.

#### 2.2.2 Extraction of AME, TeA and ATX-II

For each time point, metabolism was stopped by adding 200 µL of ice-cold (-20 °C) ACN/MeOH (50:50, *v/v*). Thereafter, the tubes were sonicated for 10 min in a water bath, then the suspensions were transferred to 1.5 mL low-binding Eppendorf tubes and centrifuged for 5 min at 10,000 × *g* at room temperature. The supernatants were transferred to 2 mL amber glass vials and stored at -80 °C until analysis.

#### 2.2.3 Stability assays

The stability of the toxin at 37 °C throughout the experiment was also confirmed using technical replicates prepared and incubated under the same conditions as the kinetic experiment. Thus, 100 µL of the treatment solution was added to 100 µL of DMSO-free medium (1:1, *v/v*) and incubated at 37 °C with 5 % CO_2_ for the entire duration of the kinetic experiment. After the incubation period, the extraction step as described above was performed and measured alongside the other kinetic samples by LC-MS/MS.

### 2.3 LC-MS/MS analysis

#### 2.3.1 Sample preparation for LC-MS/MS analysis

Following storage at −80 °C, samples were thawed at room temperature, vortexed and an aliquot was transferred to a 1.5 mL reaction tube. Samples were centrifuged at 18,400 × g for 15 min at 4 °C. Additional dilutions using LC-MS-grade MeOH/H_2_O (3:7, *v/v*) were performed if necessary to ensure that analyte concentrations remained within the respective calibration ranges for AME (0.101– 101.4 µg/L), TeA (1.2–1210 µg/L), ATX-II (0.1–100 µg/L), and ATX-I (0.8–80 µg/L).

#### 2.3.2 Quantification of mycotoxins via targeted LC-MS/MS analysis

Quantification of AME, TeA, ATX-II and ATX-I was performed using an Agilent 1290 Infinity II high-performance liquid chromatography system (Agilent, Waldbronn, Germany) coupled to a SCIEX QTrap 6500+ mass spectrometer (Sciex, Darmstadt, Germany) equipped with Turbo-V™ electrospray ionization (ESI) source.

LC-MS/MS analysis was performed based on a previously developed method (Puntscher et al., 2018). In brief, for chromatographic separation, the Supelco Ascentis Express C18 column (10 cm × 2.1 mm, 2.7 µm; Supelco, Munich, Germany) equipped with a SecurityGuard™ C18 cartridge (4 × 2.0 mm ID; Phenomenex Ltd. Deutschland, Aschaffenburg, Germany) was used. Eluent A consisted of 5 mM aqueous ammonium acetate adjusted to pH 8.6 with 25 % ammonia, while eluent B was LC-MS/MS-grade MeOH. The multi-step gradient was optimized as follows: the column was maintained at 10% eluent B for the first 1.0 min, followed by an increase to 38% over the next 0.5 min. The proportion of eluent B was then increased linearly to 40% at 6 min, 58% at 6.1 min, 61% at 7.5 min, and 85% at 9 min. This was followed by an isocratic column-purging step at 100% eluent B from 9.1 to 13 min and a 2 min re-equilibration at the initial conditions. The total run time was 15.5 min (Puntscher et al., 2018). The autosampler temperature was set to 7 °C, the column oven was maintained at 30 °C and the injection volume was 2 µL. A divert valve was utilized to direct the effluent to the waste between 0 and 1 min. The mass spectrometer was operated in negative electrospray ionization (ESI) mode, with analytes detected predominantly as their deprotonated ions. Scheduled multiple reaction monitoring (sMRM) was used for the quantitative analysis of the parent compounds and ATX-I, whereas multiple reaction monitoring (MRM) was employed for targeted metabolite analysis. The different toxins were quantified using solvent-based external calibration, with a calibration set injected approximately every 20 samples. The applied sMRM and MRM transitions and optimized parameters are summarized in **Supplemental Table 2.**

Data acquisition and preliminary evaluation were performed using SCIEX OS (version 4.0.0.8559) (SCIEX LLC, Marlborough/MA, USA). Further data processing was carried out using Skyline software (version 26.1.0.057; MacCoss Lab, Department of Genome Sciences, University of Washington, Seattle/WA, USA).

#### 2.3.3 Evaluation of method performance

Method performance was evaluated by assessing apparent recovery (RA), extraction recovery (RE) and matrix effects expressed as signal suppression/enhancement (SSE), as well as the actual concentrations of the treatment solutions. Solvent-based spiking solutions containing individual analytes (AME, TeA, ATX-II, or ATX-I) were used to fortify medium samples either before or after sample preparation. For each condition, three independent sample sets were prepared and analyzed (**Supplemental Figure 1**).

The RA was attained by comparing the detected concentrations (calculated using the analyzed calibration curve in pure solvent) against the spiked concentrations. The RE was determined by comparing the mean concentrations of AME, TeA, ATX-II, and ATX-I measured in medium samples spiked prior to extraction with those obtained from medium samples spiked after extraction. The SSE attributable to the cell culture medium was evaluated by comparing the concentrations of AME, TeA, ATX-II, and ATX-I measured in post-extraction spiked medium with those obtained from solvent-based standards at the corresponding concentration level (**Supplemental Figure 1**).

### 2.4 In vitro toxicokinetics

Michaelis–Menten parameters, including the maximum reaction velocity (Vmax) and Michaelis– Menten constant (Km), were successfully determined for AME, enabling the calculation of intrinsic clearance. In contrast, the metabolic profiles of the other toxins exhibited either very rapid or very slow turnover, precluding reliable estimation of Michaelis–Menten parameters. For each AME concentration, the first-order elimination rate constant (k_inc_, [1/min]) was obtained from the negative slope of the log-linear plot of substrate concentration (ln(cₜ/c₀)) versus time, following linear regression (Baranczewski et al., 2006)) as equation (1) where C_t_ is the AME concentration (µM) at time t (min) and C0 is the initial concentration (µM)

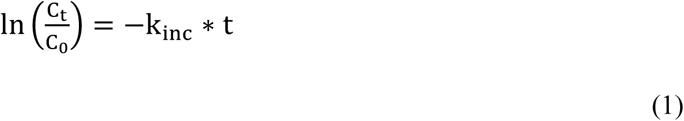

Then, V_0_, initial rate (pmol·min⁻¹·10⁶ hepatocytes⁻¹) was calculated as following where V_inc_ is the volume of incubation (µL) and N_cells, *in vitro*_ is the number of hepatocytes in the *in vitro* incubation (10^6^cells):

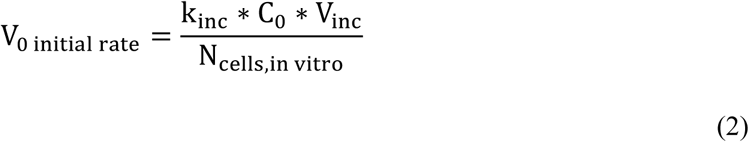

Michaelis–Menten parameters were derived using non-linear regression with GraphPad Prism 10.5.0. Thus, V_0_, initial rate (pmol·min⁻¹·10⁶ hepatocytes⁻¹) was reported in function of toxin concentration (µM), then V_max_ (pmol·min⁻¹·10⁶ hepatocytes⁻¹) and K_m_ (µM) were generated.

Intrinsic clearance (Cl_int_, [µL/min/10⁶ cells]) was calculated as following:

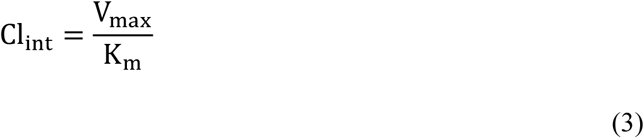

### 2.5 Statistical analysis

Significant differences among multiple groups (≥3) were evaluated by applying one-way ANOVA, followed by Fisher’s LSD as a post hoc test. Comparison between two groups was performed using the Student’s *t*-test. All statistical analyses were performed with OriginPro 2025 (Version 10.2.0.188; Academic, OriginLab Corporation, Northampton, MA, USA). All experiments were carried out in technical triplicates.

## 3. Results and discussion

### 3.1 In vitro parent compounds depletion in PRHs and PHHs

The aim of this study was to investigate the hepatic clearance of AME, TeA, and ATX-II in PRHs and PHHs and to assess their metabolic stability.

#### 3.1.1 Cytotoxicity data

ATP-based viability assays were performed to assess potential cytotoxic effects in PRHs and PHHs across five independent experiments. Cell viability was expressed relative to the corresponding solvent control (0.05% DMSO). In PRHs, mean cell viability was 86.7 ± 13.8% for AME, 74 ± 3% for TeA, and 80.1 ± 8.9% for ATX-II. In PHHs, mean cell viability was 87.7 ± 7.5% for AME, 84 ± 2% for TeA, and 103.0 ± 6.2% for ATX-II. In experiments evaluating different cell densities, mean cell viability for AME was 112.3 ± 17.1% in PRHs and 152.4 ± 39.7% in PHHs.

#### 3.1.2 Evaluation of the method performance

The apparent recovery (RA) was 104% for AME, 112% for TeA, 44% for ATX-II, and 91% for ATX-I. The extraction recovery of the sample preparation procedure showed mean recoveries of 103% for AME, 120% for TeA, 45% for ATX-II, and 79% for ATX-I (**Supplemental Figure 1**). The extraction recovery (RE) obtained for AME, TeA, and ATX-I indicate efficient and reproducible analyte extraction. The observed variability is not expected to adversely affect assay accuracy or precision and is consistent with the performance criteria outlined in the ICH M10 guideline on bioanalytical method validation (European Medicines Agency, 2023). To account for the low RA of ATX-II, measured concentrations were corrected using a recovery factor of 2.27.

Matrix effects were analyte-dependent, with signal suppression observed for TeA (85%) and ATX-II (80%), and slight signal enhancement for AME (110%) and ATX-I (116%). The lower recovery is more likely attributable to analyte-specific interactions with cellular components or other constituents of the sample matrix during sample preparation (**Supplemental Figure 1**).

The limit of detection (LOD) was determined in Skyline based on the analytical response of blank samples and the calibration curve. The detection threshold was defined as the mean blank response plus three times the standard deviation (SD) of the blank response and was subsequently converted to the corresponding analyte concentration using the fitted calibration curve. The LOD therefore represents the lowest analyte concentration, and it is expected to produce a signal distinguishable from the background response. The limit of quantification (LOQ) was estimated using a 10-SD criterion and was therefore calculated as 10/3 times the LOD (approximately 3.3 × LOD). The LOD and LLOQ for the different toxins analyzed are shown in **Supplemental Table 1.**

### 3.2 Depletion of AME in PHHs and PRHs

In stability experiments, we observed that AME was stable in the cell culture medium up to 4 h, with a negligible decrease of AME concentrations (**Supplemental Figure 2**). In contrast, in the presence of PRHs or PHHs, AME (1 µM) was completely metabolized within 30 min, with no parent compound remaining detectable, consistent with very rapid velocities (**Supplemental Figure 2**). Previously, following oral administration of a complex *Alternaria* extract administered to Sprague–Dawley rats, AME was recovered predominantly in the feces (>89%) and was not detectable in plasma, suggesting low oral bioavailability (Louro et al., 2024; Puntscher, Aichinger, et al., 2019). These findings are consistent with studies in pigs, which reported low oral bioavailability (approximately 9–15%) and high systemic clearance (approximately 13–17 L·h⁻¹·kg⁻¹) for AME, possibly indicating rapid hepatic metabolism of the absorbed fraction (den Hollander et al., 2025).

A time-dependent clearance of AME in PRHs was further evaluated at two cell densities (0.25 and 0.5 million cells/mL) using shorter incubation intervals to better characterize the clearance profile of AME (**Supplemental Figure 3**). Significant differences were found for all time points. After 5 min of incubation at 5 µM, AME concentrations decreased to 3.42 µM and 3.38 µM in the 0.5 and 0.25 million cells/mL groups, respectively, corresponding to declines of 24% and 23% relative to T0. By 10 min, concentrations had further decreased to 2.07 µM in the 0.5 million cells/mL group and 2.35 µM in the 0.25 million cells/mL group, representing overall reductions of 54 % and 47 %, respectively. By 30 min, the 0.25 million cells/mL group showed lower remaining AME concentrations than the 0.5 million cells/mL group, despite the lower cell density, 0.88 µM compared to 0.25 µM in the 0.5 and 0.25 million cells/mL groups, representing a total of 81% and 94% decline in concentrations.

The results indicate distinct species-related differences in the rate and extent of AME depletion during hepatocyte incubation. In the lower concentrations (0.25–0.75 µM), AME levels remained low in both systems, ranging from 0.09 to 0.58 µM in PRHs and from 0.14 to 0.53 µM in PHHs. At 0.25 and 0.5 µM, concentrations fell below the LOD within 10 min in both hepatocyte models **(Figure 1).** As the concentration increased to 1.5 and 3 µM, clearer differences between PRHs and PHHs became apparent, with PRHs generally showing greater reduction in AME concentration at the 10 min time point compared with PHHs. At the highest tested concentration (8 µM), this trend persisted. In PRHs, AME concentrations decreased from 7.28 µM at T0 to 5.49 µM at 10 min, whereas in PHHs, concentrations decreased from 6.19 µM to 5.87 µM **(Figure 1)**. Overall, these findings demonstrate that AME undergoes hepatocellular clearance in both rat and human hepatocytes, with PRHs exhibiting consistently higher clearance efficiency than PHHs, particularly at moderate to high concentrations.

Species-dependent differences in hepatic metabolism have been described previously for several mycotoxins. For example, Gross-Steinmeyer *et al*. (2002) reported that ochratoxin A was metabolized to several oxidative and conjugated metabolites in PRHs, whereas metabolism in PHHs was minimal, highlighting substantial interspecies differences in hepatic metabolic capacity (Gross-Steinmeyer et al., 2002). Similar observations have recently been reported for the structurally related dibenzo-α-pyrone altenuene (ALT), for which substrate depletion reached approximately 88% in rat hepatocytes compared with 57% in human hepatocytes after 4 h of incubation (Borsos et al., 2026). Likewise, Borsos *et al*. (2024) characterized the qualitative and quantitative aspects of phase I metabolism of alternariol (AOH) and AME using nicotinamide adenine dinucleotide phosphate-fortified-porcine, rat, and human liver microsomes. Then authors reported significant decreases in alternariol (AOH) after 5 min (1 μM for all three species; 10 and 20 μM for rat and human species) or 10 min (10, 20, and 50 μM for porcine liver microsomes). Researchers also noted phase I metabolism of AOH proceeded at comparable rates in rat and human liver microsomes, whereas porcine liver microsomes exhibited lower metabolic activity. In contrast, phase I metabolism of AME was more similar between human and porcine liver microsomes, while rat liver microsomes displayed a modestly higher metabolic rate. However, these findings should be interpreted with caution, as the study was limited to phase I metabolism and did not account for phase II conjugation reactions, which are known to play a major role in the biotransformation of both AOH and AME (Borsos et al., 2024).

Our findings show that AME was rapidly depleted in both primary rat and primary human hepatocytes, with complete disappearance of the parent compound within 30 min. The comparable depletion kinetics observed in both species suggest that AME undergoes efficient hepatic biotransformation irrespective of species and indicate that the enzymes responsible for its metabolism are highly active in both rat and human hepatocytes. While species-specific differences in the identity and relative abundance of AME metabolites remain to be elucidated, the relatively small differences in parent compound depletion suggest that other factors, including metabolite formation and extrahepatic metabolism, may contribute more substantially to interspecies differences in AME toxicokinetics.

#### 3.2.1 Determination of Michaelis–Menten parameters: Vmax and Km

Following LC-MS/MS analysis results, the depletion in the AME parent compound was measured between T0 and 10 min of incubation. For PRHs and PHHs, the first order elimination rate constant (*k*_inc_, [1/min]) was calculated using equation (1), and the initial velocity initial rate (V0) was determined using equation (2). The results are presented in **Table 1**.

**Table 1:** First-rate constant K_inc_ (1/min) and V_0_ (pmol·min^-1^·10^6^ hepatocytes⁻¹) initial rate calculations for primary rat (PRH) and human (PHH) hepatocytes.

| PRHs |  |  |  | PHHs |  |  |
| --- | --- | --- | --- | --- | --- | --- |
| *Alternariol monomethyl ether* (AME) ($\mu\text{M}$) | $C_0$ ($\mu\text{M}$) | first-rate constant $k_{inc}$ (1/min) | $V_0$ initial rate ($\text{pmol} \cdot \text{min}^{-1} \cdot 10^6 \text{ hepatocytes}^{-1}$) | $C_0$ ($\mu\text{M}$) | first-rate constant $k_{inc}$ (1/min) | $V_0$ initial rate ($\text{pmol} \cdot \text{min}^{-1} \cdot 10^6 \text{ Hepatocytes}^{-1}$) |
| 0.25 | 0.025 | 0.284 | 14.15 | 0.040 | 0.206 | 16.28 |
| 0.5 | 0.061 | 0.248 | 30.16 | 0.081 | 0.148 | 23.98 |
| 0.75 | 0.187 | 0.203 | 75.70 | 0.150 | 0.105 | 31.46 |
| 1.5 | 0.388 | 0.096 | 74.59 | 0.353 | 0.073 | 51.49 |
| 3 | 0.864 | 0.076 | 131.7 | 0.765 | 0.018 | 26.76 |
| 8 | 2.082 | 0.029 | 118.7 | 1.769 | 0.005 | 18.40 |

$C_0$ is the initial concentration ( $\mu\text{M}$ ) and $k_{inc}$ are expressed here as mean of $n=3$ technical replicates (one tube per point). The $V_0$ is the initial rate calculated for PRHs showed that saturation occurs at approximately 0.75 $\mu\text{M}$ of AME, whereas for PHHs, no plateau was observed. The results for PRHs appear reliable, as the R square value is 0.86, whereas the R square value for PHHs is 0.12, indicating uncertainty in this last result.

The disappearance of AME obtained with the PRHs described a Michaelis–Menten curve (**Figure 2**). The linear portion (described by four concentrations) is followed by plateau (described by two concentrations) on an increasing scale of AME concentrations. For PHHs, the results are much more complex. Indeed, the data seem to indicate that the decrease in the AME parent compound (sum of intracellular and extracellular compartments) does not follow a Michaelis–Menten curve. Thus, in **Table 1**, Michaelis–Menten parameters were not indicated because of this inaccuracy. Further research is needed to elucidate this elimination in human hepatocytes. Atypical kinetic profiles are well documented in *in vitro* metabolism studies and are not uncommon for the disappearance of parent compounds (Seibert & Tracy, 2021; Tracy, 2003).

**Figure 2.**
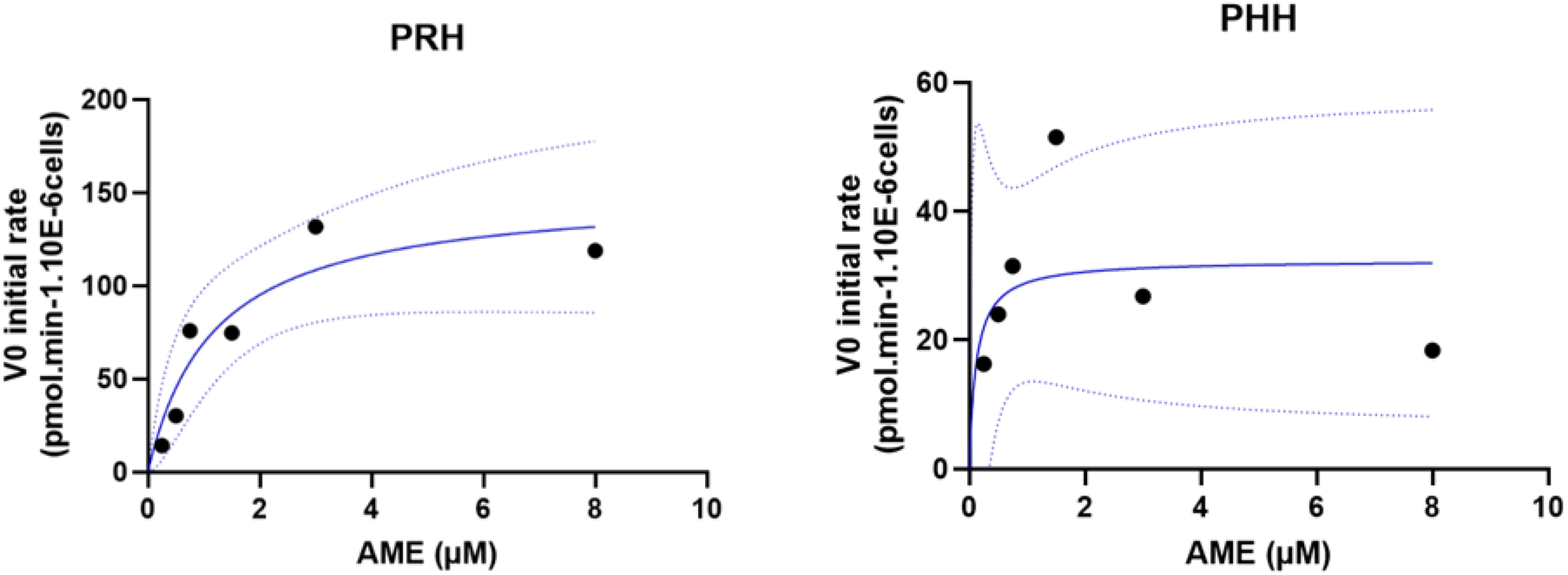
Michaelis–Menten curves for primary rat (PRH) and human (PHH) hepatocytes to determine the V_max_ (pmol·min^-1^·10^6^ hepatocytes⁻¹) and K_m_ (1/min) parameters. V_0_ initial rate (pmol·min^-1^·10^6^hepatocytes^-1^) is plotted against AME concentrations ranging from 0.25 to 8 µM. Solid blue lines represent the fitted Michaelis–Menten curves, and dotted blue lines indicate the **95% confidence intervals** of the fitted curves. A reliable Michaelis–Menten model could be established only for PRHs, whereas the PHH data did not exhibit saturable kinetics and therefore did not permit reliable estimation of kinetic parameters (Vmax and Km).

Then, intrinsic clearance was calculated using equation (3). In **Table 2**, The Vmax and Km used to calculate the Clint were reported. Intrinsic clearance differs quantitatively between rat and human hepatocytes. In fact, in addition to the higher AME velocities of disappearance observed in rats, AME metabolism in primary human hepatocytes does not appear to follow Michaelis–Menten kinetics.

**Table 2:**
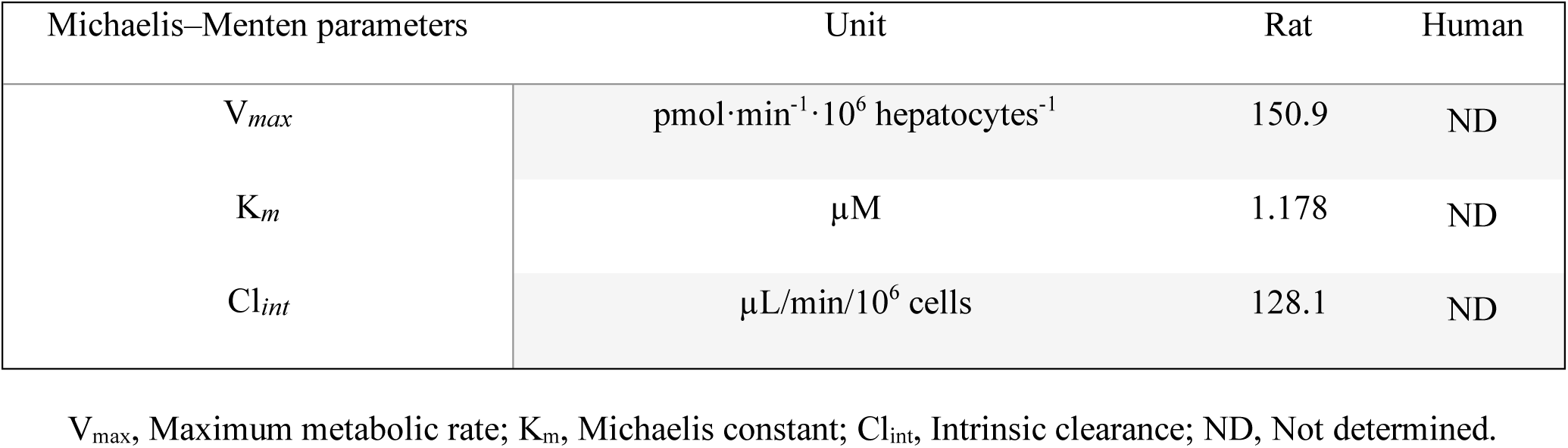
Michaelis–Menten kinetic parameters for alternariol monomethyl ether (AME) metabolism in primary rat hepatocytes (PRHs) and primary human hepatocytes (PHHs).

| Michaelis–Menten parameters | Unit | Rat | Human |
| --- | --- | --- | --- |
| $V_{max}$ | $\text{pmol} \cdot \text{min}^{-1} \cdot 10^6 \text{ hepatocytes}^{-1}$ | 150.9 | ND |
| $K_m$ | $\mu\text{M}$ | 1.178 | ND |
| $Cl_{int}$ | $\mu\text{L}/\text{min}/10^6 \text{ cells}$ | 128.1 | ND |

$V_{max}$ , Maximum metabolic rate; $K_m$ , Michaelis constant; $Cl_{int}$ , Intrinsic clearance; ND, Not determined.

### 3.3 Depletion of TeA in PHHs and PRHs

The clearance of TeA in PRHs and PHHs was assessed and remained relatively stable throughout the 4 h incubation period following treatment with the 5 µM dosing solution (**Figure 3**). At T0, the mean concentrations were 4.3 µM in PRHs and 4.1 µM in PHHs. Over 4 h, a significant difference in concentration was observed in both species, reaching 3.9 µM in rat hepatocytes and 3.7 µM in human hepatocytes, corresponding to an approximate 10% reduction in concentration. The stability control exhibited a TeA concentration of 3.9 µM, comparable to the concentrations measured following 4 h incubation with hepatocytes, indicating that TeA remained stable under the assay conditions throughout the experiment. Furthermore, the measured concentration in the treatment solutions were consistent with the target concentration of 5 µM, with values of 4.93 µM and 5.16 µM, respectively, confirming accurate preparation of the dosing solutions.

**Figure 3.**
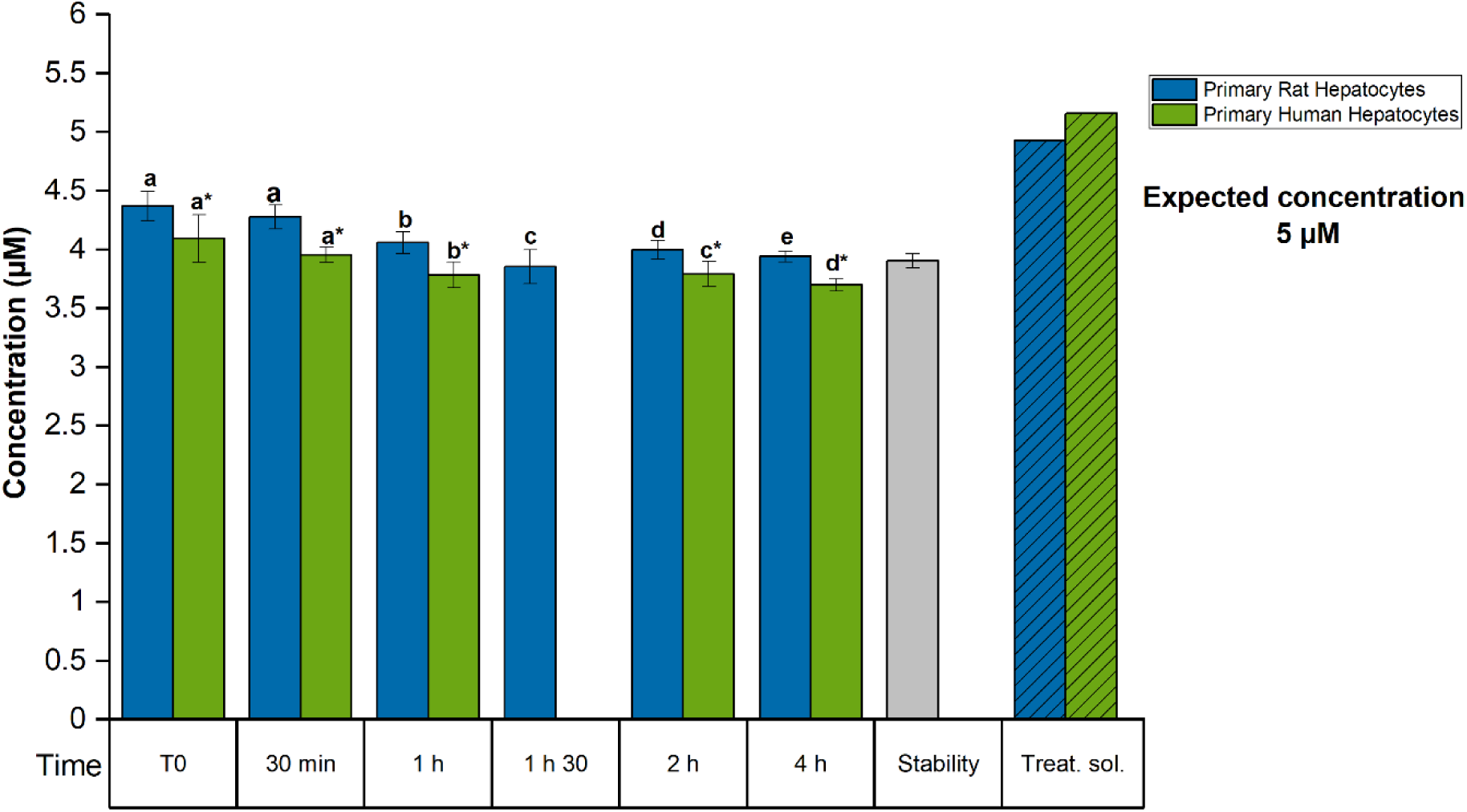
TeA concentration in PRHs and PHHs following 5 µM treatment. TeA concentrations were quantified by LC-MS/MS at multiple time points over a 4 h incubation period, alongside stability and treatment solution controls. Stability controls were prepared in the same manner as the kinetic samples by adding 100 µL of treatment solution to 100 µL of DMSO-free medium (1:1, *v/v*) and incubating at 37 °C with 5 % carbon dioxide (CO_₂_) for the full duration of the experiment in the absence of hepatocytes. Treat. sol. denotes treatment solution controls prepared by mixing 100 µL of treatment solution with 100 µL of DMSO-free medium (1:1, *v/v*), followed by immediate extraction without incubation. The TeA concentrations were quantified using liquid chromatography tandem mass spectrometry (LC-MS/MS). Results are depicted as mean ± standard deviation (SD) of three technical replicates. Statistically significant differences in TeA concentrations among multiple groups (≥3) were evaluated by applying one-way ANOVA, followed by Fisher’s LSD as a post hoc test. Comparison between two groups was performed using the Student’s *t*-test. Different letters indicate significant differences (*p* < 0.05) between T0 and subsequent time points * Correspond to PHHs. Treatment solution controls are shown without SD because only two independent treatment solutions were analyzed.

These findings are consistent with published *in vivo* toxicokinetic studies. Although *in vitro* metabolism data have previously been lacking, human volunteer studies demonstrated that approximately 90% of orally administered TeA is excreted unchanged in urine within 24 h, suggesting negligible metabolism (Asam et al., 2013). Likewise, studies in rats and mice reported that TeA is excreted largely as the unchanged parent compound, with approximately 87% of the administered dose recovered in urine within 24 h (Puntscher, Aichinger, et al., 2019; Puntscher, Hankele, et al., 2019). More recently, Visintin *et al*. (2025) reported that unmetabolized TeA remained detectable in urine for up to 13 h after ingestion, with cumulative urinary excretion over 48 h accounting for 32 ± 17% of the administered dose. In addition, TeA remained detectable in capillary blood for an average of 6.6 h, irrespective of hydrolysis treatment (Visintin et al., 2025). In the present study, no TeA metabolites were detected. Together with the published evidence, these results indicate that TeA undergoes minimal, if any, hepatic biotransformation and is eliminated predominantly via urinary excretion as the unchanged parent compound. Consistent with this interpretation, TeA exhibited negligible depletion in both PRHs and PHHs over the 4 h incubation period, precluding a reliable estimation of intrinsic clearance.

Overall, the slight decrease observed during hepatocyte incubation may therefore reflect minor nonspecific processes such as cellular uptake, limited non-enzymatic degradation rather than extensive enzymatic metabolism or adsorption to incubation materials (Aichinger et al., 2020).

### 3.4 Depletion of ATX-II in PHHs and PRHs

At T0 ATX-II concentrations were 0.03 µM in PRHs and 0.02 µM in PHHs. ATX-II concentrations decreased rapidly and were below the LOD after 30 min in both PRHs and PHHs, indicating extensive metabolism of the parent compound (**Supplemental Figure 4**). This observation was supported by the concurrent detection of ATX-I in both hepatocyte models, demonstrating partial biotransformation of ATX-II to ATX-I. In PRHs, ATX-I concentrations increased from 0.05 µM at 30 min to 0.06 µM at 2 h. In PHHs, ATX-I concentrations were consistently lower, reaching 0.03 µM at 30 min and decreasing to 0.01 µM thereafter. By the 4 h time point, neither ATX-I nor ATX-II were detectable in either hepatocyte model.

The 4 h stability control samples and treatment solutions showed measurable ATX-II concentrations of 0.1 µM, 0.14 µM and 0.12 µM, suggesting that ATX-II may exhibit partial chemical instability under the experimental conditions independent of hepatocyte-mediated metabolism (Aichinger et al., 2018). Overall, these findings demonstrate rapid clearance of ATX-II in both PRHs and PHHs, accompanied by formation of ATX-I, with metabolic conversion appearing more pronounced in PRHs than in PHHs (**Supplemental Figure 4**).

The second experiment, conducted with shorter time points (T0, 5, 10, 20, and 30 min), confirms the rapid decline of ATX-II (**Figure 3**). In fact, even after a 5 min exposure to ATX-II, the parent compound was no longer detectable; only the ATX-I metabolite, resulting from reductive de-epoxidation, could be detected and quantified. For PRH, the ATX-I metabolite was quantified at average concentrations of 51.8, 47.9, 41.4, and 40.0 nM for 5, 10, 20, and 30 min, respectively. For PHHs, the ATX-I metabolite was quantified at average values of 40.1, 26.0, 21.7, and 0 nM for exposure times of 5, 10, 20, and 30 min, respectively. These results show that the concentration of PRH stabilized after 5 min of exposure, whereas a decrease in ATX-I was observed for PHH after 5 min of exposure (**Figure 4**).

**Figure 4.**
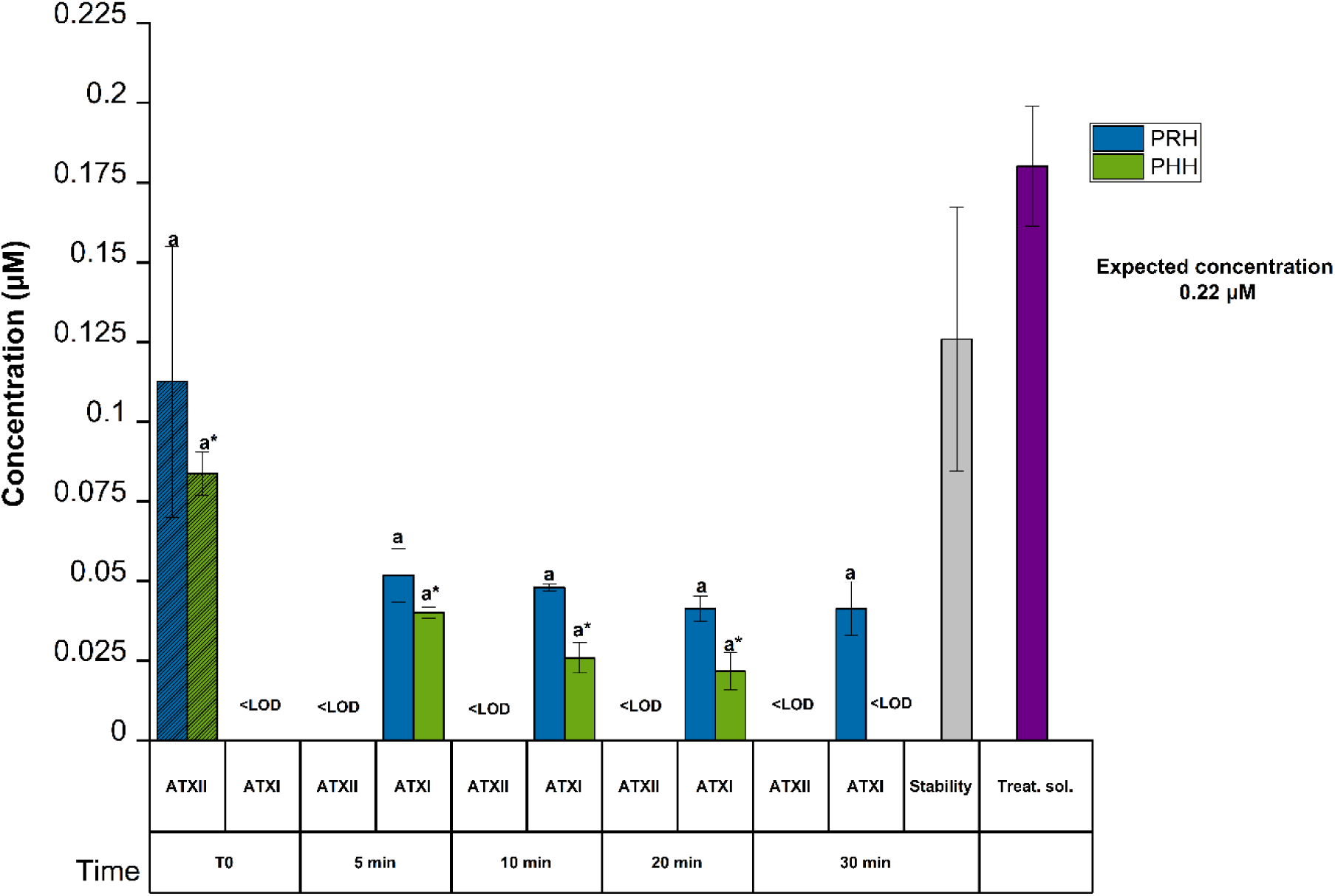
The concentrations of altertoxin II (ATX-II) and altertoxin I (ATX-I) in primary rat (PRH) and human (PHH) hepatocytes following 0.22 µM treatment. Stability controls were prepared in the same manner as the kinetic samples by adding 100 µL of treatment solution to 100 µL of DMSO-free medium (1:1, *v/v*) and incubating at 37 °C with 5 % carbon dioxide (CO_₂_) for the full duration of the experiment in the absence of hepatocytes. Treat. sol. denotes treatment solution controls prepared by mixing 100 µL of treatment solution with 100 µL of DMSO-free medium (1:1, *v/v*), followed by immediate extraction without incubation. ATX-II and ATX-I concentrations were quantified by liquid chromatography tandem mass spectrometry (LC-MS/MS) at multiple points over a 30 min incubation period. Results are depicted as mean ± standard deviation (SD) of three technical replicates. Statistically significant differences in ATX-II/ATX-I concentrations among multiple groups (≥3) were evaluated by applying one-way ANOVA, followed by Fisher’s LSD as a post hoc test. Comparison between two groups was performed using the Student’s *t*-test. Different letters indicate significant differences (*p* < 0.05) between T0 and 10 min. * Corresponds to PHHs.

These findings support previous observations by Fleck *et al*. (2014), who demonstrated that, in Caco-2 and HepG2 cells, ATX-II was rapidly converted to ATX-I, suggesting that the reduction of epoxides is a major detoxification pathway (Fleck et al., 2014). Although ATX-I is generally regarded as a detoxification product, the rapid conversion observed in the present study is toxicologically relevant because the parent compound ATX-II is capable of forming covalent guanine DNA adducts via its epoxide functionality (Soukup et al., 2020). Thus, rapid disappearance of ATX-II from hepatocyte incubations may represent an important cellular defense mechanism. Supporting this interpretation, Call *et al*. (2026) also reported a progressive decline in ATX-II concentrations during standardized *in vitro* digestion of tomato purée spiked with a complex *Alternaria* mycotoxin extract, with only 4–12% of the initial ATX-II remaining after completion of the intestinal phase. Although ATX-I formation was not detected under these digestion conditions, the marked loss of ATX-II further highlights its intrinsic instability and susceptibility to biotransformation. Moreover, *in vivo* studies in rats demonstrated that oral administration of either mixed *Alternaria* extracts or isolated ATX-II resulted in the systemic detection of ATX-I in plasma, urine, and feces, whereas ATX-II was not detected in all matrices, providing strong evidence that reductive de-epoxidation occurs *in vivo* (Puntscher, Aichinger, et al., 2019; Puntscher, Hankele, et al., 2019). Collectively, these findings, together with the present results, support the rapid conversion of ATX-II to ATX-I as a major detoxification pathway across different biological systems. The extensive depletion of the parent compound under the present experimental conditions also precluded reliable estimation of intrinsic clearance.

### 3.5 Tentative targeted analysis of selected AME metabolites via LC-MS/MS

Tentative targeted LC–MS/MS analysis of metabolites demonstrated that AME underwent both phase I and phase II biotransformation in primary rat and human hepatocytes. Phase I metabolism resulted in the formation of O-demethylated and hydroxylated metabolites, whereas phase II metabolism was characterized by glucuronide and sulfate conjugation (**Supplemental Figures 5 and 6**).

The glucuronidation observed in the present study is in agreement with previous research. More recently, Borsos et *al.* (2026) reported that the targeted approach found that the structurally similar ALT exclusively undergoes conjugation reactions via glucuronidation in both PRH and PHH (Borsos et al., 2026). Previously, glucuronidation assays using rat, pig, and human hepatic and intestinal microsomes identified AME-3-*O*-glucuronide as the major conjugation product, whereas AME-7-*O*-glucuronide was formed to a lesser extent (Pfeiffer et al., 2009). In a study where the incubation of AME with Caco-2 cells was carried out, results yielded AME-3-*O*-glucuronide, AME-7-*O*-glucuronide, and AME-3-*O*-sulfate after 2 h (Burkhardt et al., 2009).

Additionally, the oxidative metabolism of AME has been investigated in several *in vitro* systems. Early studies using rat liver S9 fractions supplemented with an NADPH-regenerating system demonstrated extensive phase I metabolism, including *O*-demethylation to alternariol (AOH) (Pollock et al., 1982). Subsequently, studies employing rat, pig, and human hepatic microsomes identified six hydroxylated metabolites (2-, 4-, 8-, and 10-OH-AME, dihydroxy-AME, and methylhydroxy-AME) in addition to AOH, revealing pronounced species differences in oxidative metabolism (Pfeiffer et al., 2007).

The metabolite profiles indicated both conserved and species-dependent patterns of AME biotransformation in rat and human hepatocytes (**Supplemental Figures 5 and 6**). Both systems generated metabolites consistent with O-demethylation, hydroxylation, glucuronidation, and sulfation, supporting broadly conserved phase I and phase II metabolic pathways. However, marked species differences were observed at intermediate AME concentrations. At 1.5 µM, rat hepatocytes showed substantially greater formation of several metabolites, particularly the AME sulfate (AME-3-S), AOH sulfate (AOH-3-S-1) and hydroxylated AOH (OH-AOH), whereas the corresponding metabolite responses were considerably lower in human hepatocytes. In contrast, OH-AOH became the predominant metabolite in both species at higher AME concentrations. Formation of AOH and the second hydroxylated AOH metabolite also increased at the highest concentration and reached comparable levels in rat and human hepatocytes. Overall, these profiles suggest a greater capacity for AME biotransformation in rat hepatocytes at low-to-intermediate substrate concentrations, particularly through sulfation and hydroxylation, while the interspecies differences became less pronounced at 8 µM.

The absence of authentic reference standards for AME metabolites limited metabolite quantification to a semi-quantitative assessment based on LC-MS/MS peak areas. This approach assumes comparable ionization efficiencies among structurally related AME-derived analytes analyzed under identical chromatographic, matrix, and instrumental conditions. However, ionization efficiencies may vary by several orders of magnitude, representing an inherent limitation of this analytical approach (Liigand et al., 2018). Consequently, the reported metabolite abundances should be interpreted as relative rather than absolute. Despite this limitation, the present study provides novel insights into the metabolism of the comparatively understudied mycotoxin AME in physiologically relevant primary hepatocyte models, which retain the coordinated activity of both phase I and phase II drug-metabolizing enzymes. These semi-quantitative metabolic profiles provide a valuable basis for prioritizing metabolites for future structural elucidation and toxicological characterization.

## 4. Conclusion

The current study offers a quantitative characterization of the *in vitro* hepatic clearance of the understudied *Alternaria* toxins AME, TeA and ATX-II in primary rat and human hepatocytes. The findings demonstrate pronounced compound- and species-dependent differences in hepatic biotransformation. AME underwent rapid hepatic clearance in both hepatocyte models, with consistently higher clearance rates observed in rat hepatocytes than in human hepatocytes. Moreover, Michaelis– Menten kinetics could be established for rat hepatocytes, indicating species-specific differences in the enzymatic processes governing AME metabolism, marking a notable contribution to the field. In contrast, TeA exhibited negligible hepatic clearance in both species, suggesting that hepatic metabolism plays only a minor role in its elimination. ATX-II was rapidly depleted in both hepatocyte models and was accompanied by transient formation of ATX-I, supporting rapid biotransformation of ATX-II.

Tentative analysis of the occurring AME metabolites revealed that AME undergoes phase I and phase II metabolism. These findings contribute to closing a critical knowledge gap in the toxicokinetics of this understudied *Alternaria* toxin and provide a foundation for future risk assessment efforts.

Overall, these findings improve current understanding of the toxicokinetic behavior of these *Alternaria* mycotoxins and highlight the importance of considering species differences during the process, which often begins with rodents and leads to the characterization of the risk to humans. The intrinsic clearance parameters and metabolic profiles generated in this study provide valuable input for physiologically based toxicokinetic modelling and human health risk assessment. Future studies should further characterize the identification of metabolites and associated enzymes involved in the metabolism of these toxins and evaluate the toxicological relevance of the resulting biotransformation products.

## Notes

### Competing Interest Statement

The authors have declared no competing interest.

### Summary of Updates

Results updated; Figures revised; author affiliations updated; Supplemental files updated.

### Funding

- Horizon Europe: 101057014

## References

Aichinger, G., Del Favero, G., Warth, B., & Marko, D. (2021). Alternaria toxins—Still emerging? Comprehensive Reviews in Food Science and Food Safety, 20(5), 4390–4406. 10.1111/1541-4337.12803

Aichinger, G., Puntscher, H., Beisl, J., Kütt, M.-L., Warth, B., & Marko, D. (2018). Delphinidin protects colon carcinoma cells against the genotoxic effects of the mycotoxin altertoxin II. Toxicology Letters, 284, 136–142. 10.1016/j.toxlet.2017.12.002

Aichinger, G., Živná, N., Varga, E., Crudo, F., Warth, B., & Marko, D. (2020). Microfiltration results in the loss of analytes and affects the in vitro genotoxicity of a complex mixture of Alternaria toxins. Mycotoxin Research, 36(4), 399–408. 10.1007/s12550-020-00405-9

Asam, S., Habler, K., & Rychlik, M. (2013). Determination of tenuazonic acid in human urine by means of a stable isotope dilution assay. Analytical and Bioanalytical Chemistry, 405(12), 4149–4158. 10.1007/s00216-013-6793-5

Baranczewski, P., Stańczak, A., Sundberg, K., Svensson, R., Wallin, A., Jansson, J., Garberg, P., & Postlind, H. (2006). Introduction to in vitro estimation of metabolic stability and drug interactions of new chemical entities in drug discovery and development. Pharmacological Reports: PR, 58(4), 453–472.

Borsos, E., Gendre, C., Mahdjoub, M., Varga, E., Dubreil, E., Henri, J., Hegarat, L. L., & Marko, D. (2026). Comparative metabolism of the Alternaria toxins altenuene and tentoxin in rat and human primary hepatocytes (p. 2026.05.11.724251). bioRxiv. 10.64898/2026.05.11.724251

Borsos, E., Varga, E., Aichinger, G., & Marko, D. (2024). Unraveling Interspecies Differences in the Phase I Hepatic Metabolism of Alternariol and Alternariol Monomethyl Ether: Closing Data Gaps for a Comprehensive Risk Assessment. Chemical Research in Toxicology, 37(8), 1356– 1363. 10.1021/acs.chemrestox.4c00095

Burkhardt, B., Pfeiffer, E., & Metzler, M. (2009). Absorption and metabolism of the mycotoxins alternariol and alternariol-9-methyl ether in Caco-2 cells in vitro. Mycotoxin Research, 25(3), 149–157. 10.1007/s12550-009-0022-2

Call, F., Crudo, F., & Marko, D. (2026). Impact of *in vitro* digestion on the bioaccessibility, genotoxicity and mutagenicity of mycotoxins in a complex *Alternaria* extract. Food Chemistry: X, 33, 103481. 10.1016/j.fochx.2026.103481

Crudo, F., Varga, E., Aichinger, G., Galaverna, G., Marko, D., Dall’Asta, C., & Dellafiora, L. (2019). Co-Occurrence and Combinatory Effects of Alternaria Mycotoxins and Other Xenobiotics of Food Origin: Current Scenario and Future Perspectives. Toxins, 11(11), 640. 10.3390/toxins11110640

den Hollander, D., De Baere, S., Holvoet, C., Devreese, M., Antonissen, G., Martens, A., Demeyere, K., Audenaert, K., Meyer, E., & Croubels, S. (2025). Absolute oral bioavailability, quantitative toxicokinetics and metabolite profiling of alternariol and alternariol monomethyl ether in pigs. Archives of Toxicology, 99(7), 2801–2817. 10.1007/s00204-025-04050-y

EFSA, Arcella, dAVIDE, Eskola, M., & Ruiz, J. A. G. (2016). Dietary exposure assessment to Alternaria toxins in the European population. EFSA Journal, 14(12), e04654. 10.2903/j.efsa.2016.4654

European Commission. (2023). Commission Regulation (EU) 2023/915 of 25 April 2023 on maximum levels for certain contaminants in food and repealing Regulation (EC) No 1881/2006 (Text with EEA relevance). http://data.europa.eu/eli/reg/2023/915/oj

European Medicines Agency. (2023). ICH M10 on bioanalytical method validation—Scientific guideline | European Medicines Agency (EMA). https://www.ema.europa.eu/en/ich-m10-bioanalytical-method-validation-scientific-guideline

Fleck, S. C., Pfeiffer, E., Podlech, J., & Metzler, M. (2014). Epoxide Reduction to an Alcohol: A Novel Metabolic Pathway for Perylene Quinone-Type Alternaria Mycotoxins in Mammalian Cells. Chemical Research in Toxicology, 27(2), 247–253. 10.1021/tx400366w

Gross-Steinmeyer, K., Weymann, J., Hege, H.-G., & Metzler, M. (2002). Metabolism and Lack of DNA Reactivity of the Mycotoxin Ochratoxin A in Cultured Rat and Human Primary Hepatocytes. Journal of Agricultural and Food Chemistry, 50(4), 938–945. 10.1021/jf0111817

Li, A. P. (2014). In Vitro Human Hepatocyte-Based Experimental Systems for the Evaluation of Human Drug Metabolism, Drug-Drug Interactions, and Drug Toxicity in Drug Development. Current Topics in Medicinal Chemistry, 14(11), 1325–1338. 10.2174/1568026614666140506114411

Liigand, P., Liigand, J., Cuyckens, F., Vreeken, R. J., & Kruve, A. (2018). Ionisation efficiencies can be predicted in complicated biological matrices: A proof of concept. Analytica Chimica Acta, 1032, 68–74. 10.1016/j.aca.2018.05.072

Louro, H., Vettorazzi, A., López de Cerain, A., Spyropoulou, A., Solhaug, A., Straumfors, A., Behr, A.-C., Mertens, B., Žegura, B., Fæste, C. K., Ndiaye, D., Spilioti, E., Varga, E., Dubreil, E., Borsos, E., Crudo, F., Eriksen, G. S., Snapkow, I., Henri, J., … Marko, D. (2024). Hazard characterization of Alternaria toxins to identify data gaps and improve risk assessment for human health. Archives of Toxicology, 98(2), 425–469. 10.1007/s00204-023-03636-8

Mahmoud, M. M., Abdel-Razek, A. S., Soliman, H. S. M., Ponomareva, L. V., Thorson, J. S., Shaaban, K. A., & Shaaban, M. (2022). Diverse polyketides from the marine endophytic *Alternaria* sp. LV52: Structure determination and cytotoxic activities. Biotechnology Reports, 33, e00628. 10.1016/j.btre.2021.e00628

Marin, S., Ramos, A. J., Cano-Sancho, G., & Sanchis, V. (2013). Mycotoxins: Occurrence, toxicology, and exposure assessment. Food and Chemical Toxicology, 60, 218–237. 10.1016/j.fct.2013.07.047

Mikula, H., Skrinjar, P., Sohr, B., Ellmer, D., Hametner, C., & Fröhlich, J. (2013). Total synthesis of masked *Alternaria* mycotoxins—Sulfates and glucosides of alternariol (AOH) and alternariol-9-methyl ether (AME). Tetrahedron, 69(48), 10322–10330. 10.1016/j.tet.2013.10.008

Pfeiffer, E., Eschbach, S., & Metzler, M. (2007). Alternaria toxins: DNA strand-breaking activity in mammalian cellsin vitro. Mycotoxin Research, 23(3), 152–157. 10.1007/BF02951512

Pfeiffer, E., Herrmann, C., Altemöller, M., Podlech, J., & Metzler, M. (2009). Oxidative in vitro metabolism of the Alternaria toxins altenuene and isoaltenuene. Molecular Nutrition & Food Research, 53(4), 452–459. 10.1002/mnfr.200700501

Pollock, G. A., DiSabatino, C. E., Heimsch, R. C., & Coulombe, R. A. (1982). The distribution, elimination, and metabolism of 14C-alternariol monomethyl ether. Journal of Environmental Science and Health, Part B, 17(2), 109–124. 10.1080/03601238209372306

Puntscher, H., Aichinger, G., Grabher, S., Attakpah, E., Krüger, F., Tillmann, K., Motschnig, T., Hohenbichler, J., Braun, D., Plasenzotti, R., Pahlke, G., Höger, H., Marko, D., & Warth, B. (2019). Bioavailability, metabolism, and excretion of a complex Alternaria culture extract versus altertoxin II: A comparative study in rats. Archives of Toxicology, 93(11), 3153–3167. 10.1007/s00204-019-02575-7

Puntscher, H., Hankele, S., Tillmann, K., Attakpah, E., Braun, D., Kütt, M.-L., Del Favero, G., Aichinger, G., Pahlke, G., Höger, H., Marko, D., & Warth, B. (2019). First insights into *Alternaria* multi-toxin *in vivo* metabolism. Toxicology Letters, 301, 168–178. 10.1016/j.toxlet.2018.10.006

Puntscher, H., Kütt, M.-L., Skrinjar, P., Mikula, H., Podlech, J., Fröhlich, J., Marko, D., & Warth, B. (2018). Tracking emerging mycotoxins in food: Development of an LC-MS/MS method for free and modified Alternaria toxins. Analytical and Bioanalytical Chemistry, 410(18), 4481–4494. 10.1007/s00216-018-1105-8

Seibert, E., & Tracy, T. S. (2021). Fundamentals of Enzyme Kinetics: Michaelis-Menten and Non-Michaelis-Type (Atypical) Enzyme Kinetics. Methods in Molecular Biology, 2342, 3–27. 10.1007/978-1-0716-1554-6_1

Soukup, S. T., Fleck, S. C., Pfeiffer, E., Podlech, J., Kulling, S. E., & Metzler, M. (2020). DNA reactivity of altertoxin II: Identification of two covalent guanine adducts formed under cell-free conditions. Toxicology Letters, 331, 75–81. 10.1016/j.toxlet.2020.05.018

Tölgyesi, Á., Stroka, J., Tamosiunas, V., & Zwickel, T. (2015a). Simultaneous analysis of Alternaria toxins and citrinin in tomato: An optimised method using liquid chromatography-tandem mass spectrometry. Food Additives & Contaminants: Part A, 32(9), 1512–1522. 10.1080/19440049.2015.1072644

Tracy, T. S. (2003). Atypical Enzyme Kinetics: Their Effect on In Vitro-In Vivo Pharmacokinetic Predictions and Drug Interactions. Current Drug Metabolism, 4(5), 341–346. 10.2174/1389200033489280

Visintin, L., Lu, E.-H., Lin, H.-C., Bader, Y., Nguyen, T. N., Michailidis, T. M., De Saeger, S., Chiu, W. A., & De Boevre, M. (2025). Derivation of human toxicokinetic parameters and internal threshold of toxicological concern for tenuazonic acid through a human intervention trial and hierarchical Bayesian population modeling. Journal of Exposure Science & Environmental Epidemiology, 35(4), 632–643. 10.1038/s41370-025-00746-6

