## Supplementary Material for "Species-dependent clearance of alternariol monomethyl ether, tenuazonic acid and altertoxin II in rat and human primary hepatocytes"

### 1. Materials and methods

1.1 Isolation, purification, and structural confirmation of ATX-II from that extract

Altertoxin-II (ATX II) was isolated from the *Alternaria* culture extract using flash chromatography conducted on an Interchim puriFlash 4250 system (Montluçon, France), endowed with Evaporative Light Scattering Detector, Photodiode Array Detector and a fraction collector, controlled by Interchim Software. For the fractionation of the extract, an Interchim PuriFlash C18-HQ column (15 µm, 12 g) was used. The mobile phases consisted of water (A) and acetonitrile (B) applying the following gradient: 5% B at 0 min, 5%-30% B in 30 min, 30% B for 30 min, 30%-98% B in 40 min, 98% B for 10 min. The flow rate was set to 6 mL/min, and 6 mL were collected per tube. In total, 200 mg extract were separated in two chromatographic runs. Thin-layer chromatography (TLC) analysis was performed, and tubes containing ATX-II were pooled into one fraction, yielding 6.22 mg of pure ATX-II.

TLC was carried out on Merck silica gel 60 PF254 plates. The mobile phase consisted of ethyl acetate and was analyzed under $UV_{254}$, and at visible light after derivatization with vanillin (1% in methanol)/sulfuric acid (5% in methanol).

Nuclear magnetic resonance (NMR) experiments were conducted on a spectrometer consisting of a Bruker Ascend 500 MHz magnet (Bruker, Billerica, MA, USA) equipped with a 5 mm triple-resonance Prodigy CryoProbe, an Avance NEO console and a SampleJet automated sample changer. Samples were measured at 298 K in deuterated methanol (Sigma-Aldrich, Co., St. Louis, MO, USA) referenced to the residual nondeuterated solvent signals ($\delta_H$ 3.31 ppm; $\delta_C$ 49.0 ppm). The resonance frequencies were 500.19 MHz and 125.77 MHz for $^1H$ NMR and $^{13}C$ NMR, respectively. Standard 1D ($^1H$, and $^{13}C$) and gradient-enhanced (ge) 2D experiments, including COSY, HSQC, and HMBC, were used as supplied by the manufacturer. The NMR spectroscopic data of ATX-II were identical to those reported in literature (Mahmoud et al., 2022).

**Tables and figures**

**Supplemental Table 1**: LOD and LOQ values for investigated mycotoxins in undiluted samples in the measurement solution

| Mycotoxin | LOD (µM) | LOQ (µM) | LOD (µg/L) | LOQ (µg/L) |
|---|---|---|---|---|
| AME | 0.006 | 0.020 | 1.63 | 5.44 |
| TeA | 0.023 | 0.076 | 4.54 | 14.99 |
| ATX-II | 0.002* | 0.011* | 0.70* | 3.85* |
| ATX-I | 0.004 | 0.013 | 1.41 | 4.58 |

AME, alternariol monomethyl ether; TeA, tenuazonic acid; ATX-II, altertoxin II; ATX-I, altertoxin I; LOD, limit of detection; LLOQ, lower limit of quantification.

*For ATX-II, LOD and LLOQ values were corrected using a recovery factor of 2.27.

**Supplemental Table 2:** Mass spectrometric parameters of the analytes measured via LC-MS/MS in negative ionization mode. An entrance potential of -10 V and a dwell time of 10 ms was universally set when operating in MRM mode. All transitions were in the same method. Quantifier transitions are marked bold where applicable.

| **Molecule** | **Precursor ion (*m/z*)** | **Product ion (*m/z*)** | **DP (V)** | **CE (V)** | **CXP (V)** | **Transition** |
|---|---|---|---|---|---|---|
| Alternariol (AOH) | 256.9 | 212.9 | -12 | -32 | -15 | Qualifier |
| | | 214.9 | -120 | -36 | -13 | Qualifier |
| | | **211.9** | **-120** | **-40** | **-13** | **Quantifier** |
| Alternariol monomethyl ether (AME) | 270.9 | 182.9 | -125 | -54 | -13 | Qualifier |
| | | 227.9 | -125 | -40 | -15 | Qualifier |
| | | **255.9** | **-125** | **-30** | **-17** | **Quantifier** |
| Alternariol monomethyl ether glucuronide (AME-GlcA) | 447.1 | 271.1 | -20 | -30 | -11 | Qualifier |
| | | **256** | **-20** | **-60** | **-11** | **Quantifier** |
| Alternariol monomethyl ether 3-sulfate (AME-3-S) | 351.0 | 271.1 | -60 | -50 | -10 | Qualifier |
| | | **256.0** | **-60** | **-50** | **-10** | **Quantifier** |
| Hydroxylated alternariol (OH-AOH) | 273 | 214.2 | -60 | -40 | -11 | Qualifier |
| | | **258.0** | **-60** | **-30** | **-11** | **Quantifier** |
| Hydroxylated alternariol monomethyl ether (OH-AME) | 287.1 | 228.0 | -40 | -30 | -11 | **Quantifier** |
| | | 272.1 | -40 | -30 | -11 | Qualifier |
| Altertoxin II (ATX-II) | 348.8 | 256.0 | -20 | -60 | -11 | Qualifier |
| | | 312.9 | -100 | -44 | -19 | Qualifier |
| | | **331.0** | **-100** | **-32** | **-19** | **Quantifier** |
| Altertoxin I (ATX-I) | 350.9 | 263.0 | -90 | -22 | -19 | Qualifier |
| | | 314.9 | -90 | -44 | -17 | Qualifier |
| | | **332.9** | **-90** | **-16** | **-21** | **Quantifier** |
| Tenuazonic acid (TeA) | 195.9 | 111.9 | -75 | -26 | -15 | Qualifier |
| | | 138.9 | -75 | -32 | -13 | **Quantifier** |

DP, declustering potential; CE, collision energy; CXP, collision cell exit potential

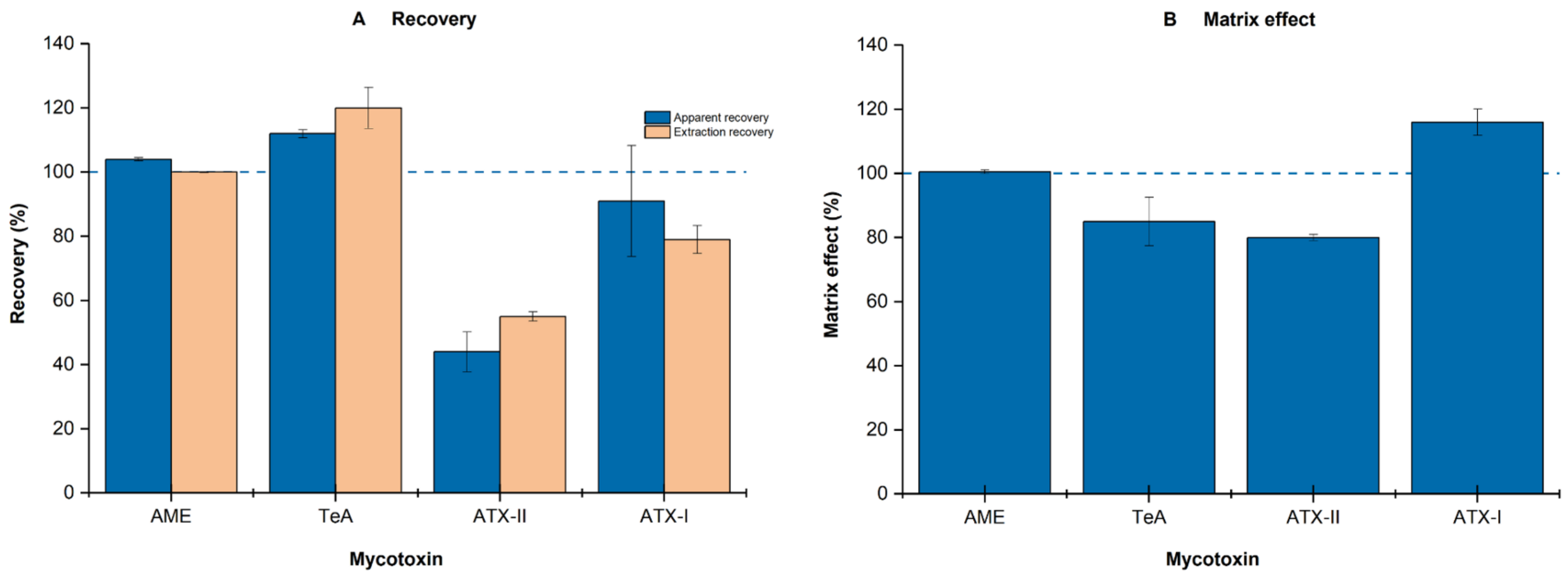


**Supplemental Figure 1.** (A) Apparent recovery and extraction recovery of alternariol monomethyl ether (AME), tenuazonic acid (TeA), altertoxin II (ATX-II), and altertoxin I (ATX-I). (B) Matrix effects determined for the four analytes. The dashed horizontal line represents 100% recovery or matrix response. Matrix-effect values below 100% indicate signal suppression, whereas values above 100% indicate signal enhancement.

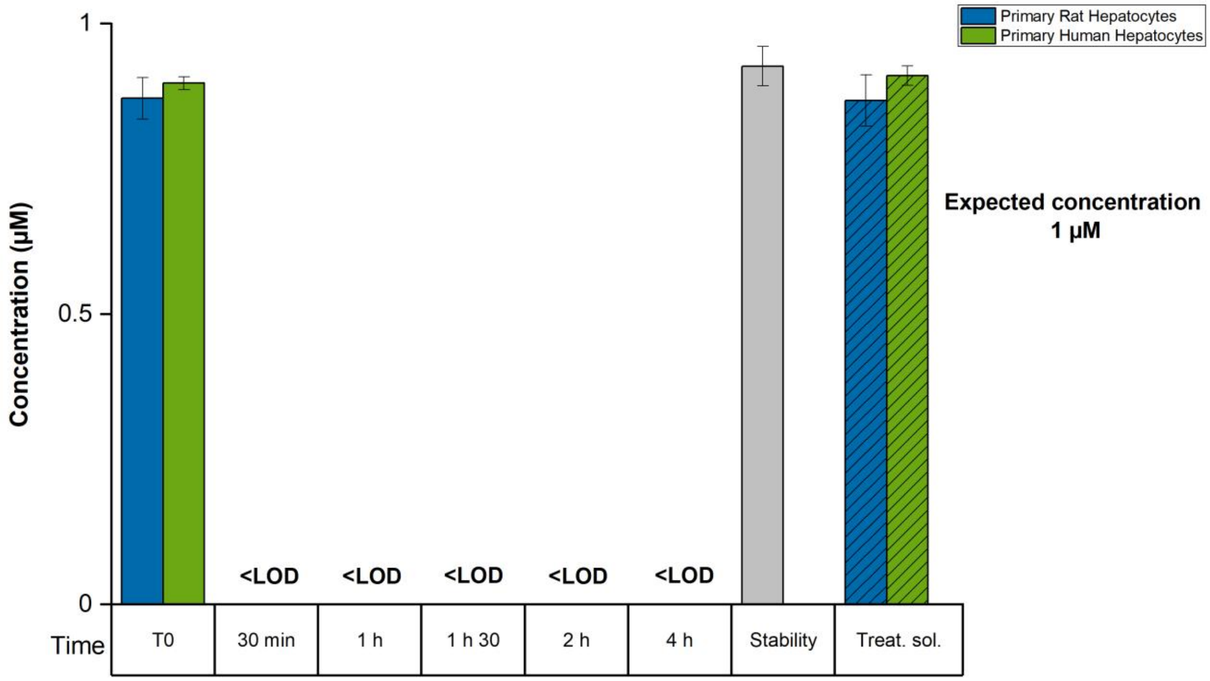


**Supplemental Figure 2**. The concentration of alternariol monomethyl ether (AME) in primary rat (PRH) and human (PHH) hepatocytes following 1 µM treatment. Stability controls were prepared in the same manner as the kinetic samples by adding 100 µL of treatment solution to 100 µL of DMSO-free medium (1:1, *v/v*) and incubating at 37 °C with 5 % carbon dioxide ($CO_2$) for the full duration of the experiment in the absence of hepatocytes. Treat. sol. denotes treatment solution controls prepared by mixing 100 µL of treatment solution with 100 µL of DMSO-free medium (1:1, *v/v*), followed by immediate extraction without incubation. AME concentrations were quantified by liquid chromatography–tandem mass spectrometry (LC-MS/MS) at multiple time points over a 4 h incubation period, alongside the stability and treatment solution controls. Results are depicted as mean ± standard deviation (SD) of three technical replicates. Treatment solution controls are shown without SD because only two independent treatment solutions were analyzed.

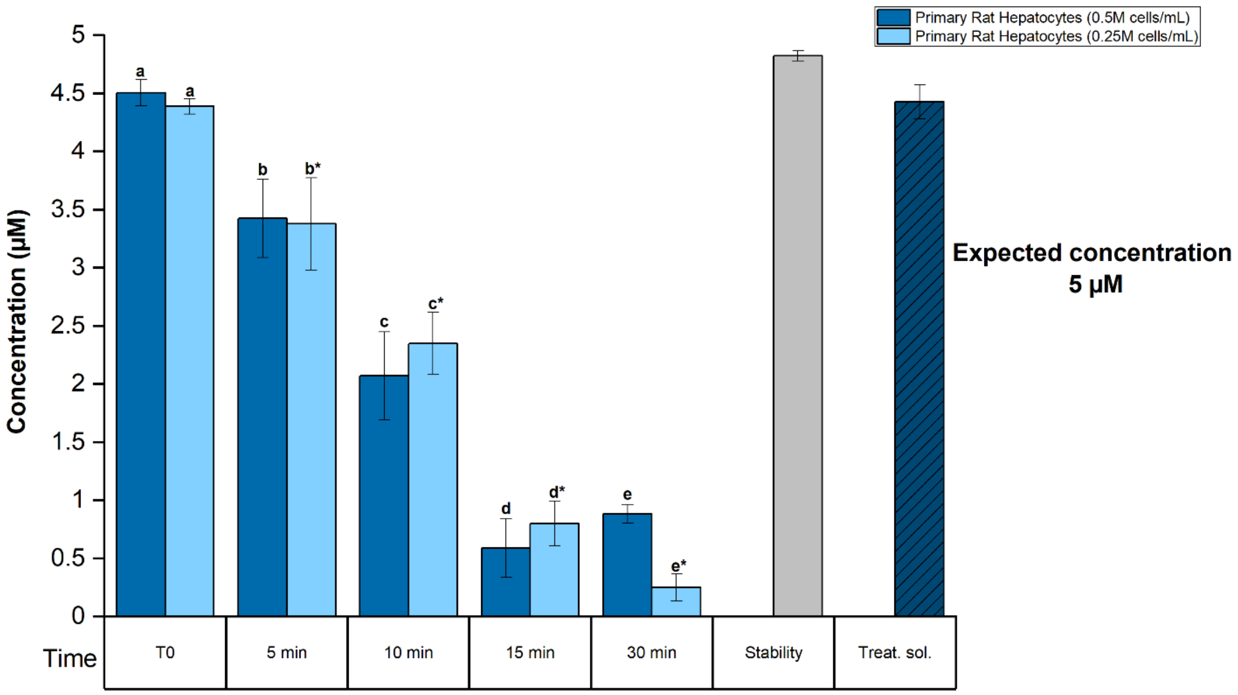


**Supplemental Figure 3.** Time-dependent clearance of alternariol monomethyl ether (AME) in primary rat hepatocytes (PRH) at two cell densities (0.25 and 0.5 million cells/mL) following 5 µM treatment. Stability controls were prepared in the same manner as the kinetic samples by adding 100 µL of treatment solution to 100 µL of DMSO-free medium (1:1, *v/v*) and incubating at 37 °C with 5 % carbon dioxide ($CO_2$) for the full duration of the experiment in the absence of hepatocytes. Treat. sol. denotes treatment solution controls prepared by mixing 100 µL of treatment solution with 100 µL of DMSO-free medium (1:1, *v/v*), followed by immediate extraction without incubation. AME concentrations were quantified by liquid chromatography–tandem mass spectrometry (LC-MS/MS) at multiple time points over a 30 min incubation period, alongside the stability and treatment solution controls. Results are depicted as mean ± standard deviation (SD) of three technical replicates. Statistically significant differences in AME concentrations among multiple groups (≥3) were evaluated by applying one-way ANOVA, followed by Fisher's LSD as a post hoc test. Comparison between two groups was performed using the Student's *t*-test. Different letters indicate significant differences ($p < 0.05$) between T0 and subsequent time points. *Corresponds to primary human hepatocytes (PHHs) (0.25M cells/mL).

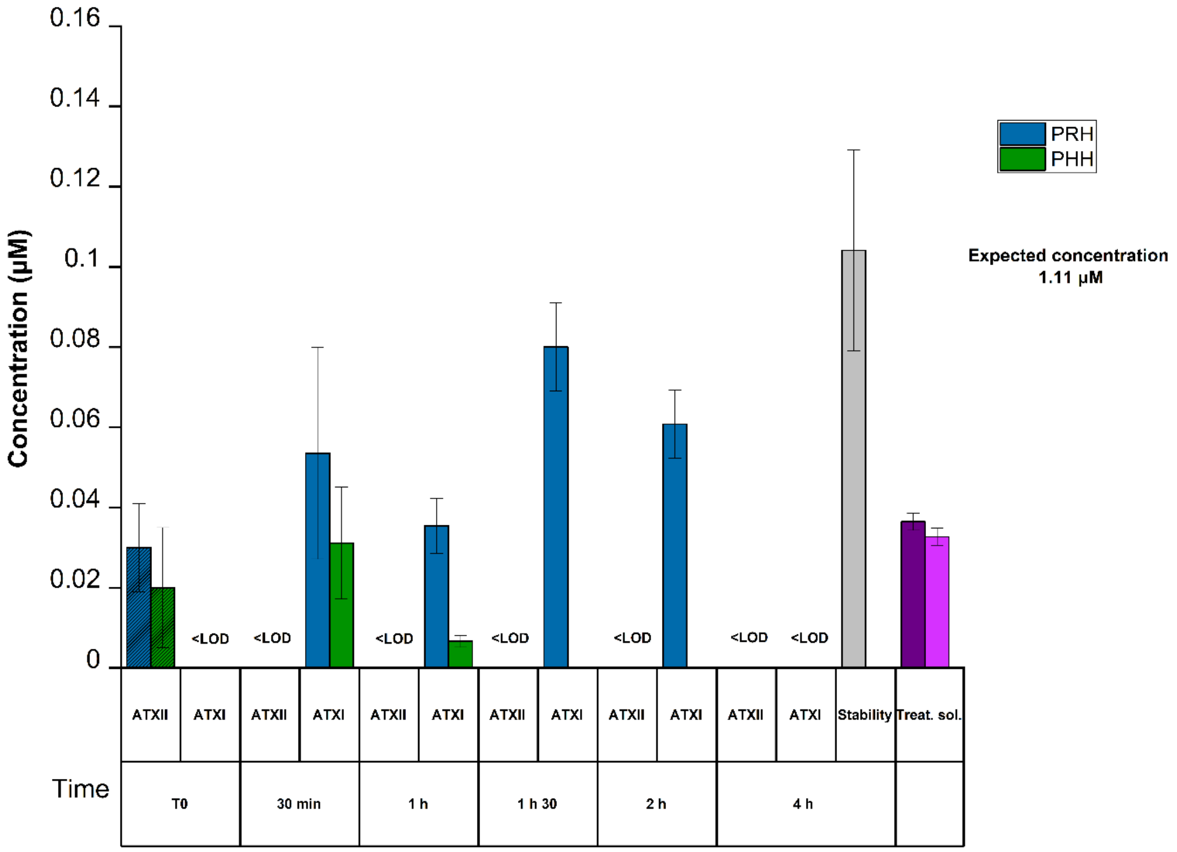


**Supplemental Figure 4**. Concentrations of alteroxin II (ATX-II) and alteroxin I (ATX-I) in primary rat (PRH) and human (PHH) hepatocytes following 1.11 µM treatment. Stability controls were prepared in the same manner as the kinetic samples by adding 100 µL of treatment solution to 100 µL of DMSO-free medium (1:1, *v/v*) and incubating at 37 °C with 5 % carbon dioxide ($CO_2$) for the full duration of the experiment in the absence of hepatocytes. Treat. sol. denotes treatment solution controls prepared by mixing 100 µL of treatment solution with 100 µL of DMSO-free medium (1:1, *v/v*), followed by immediate extraction without incubation. ATX-II and ATX-I concentrations were quantified by liquid chromatography–tandem mass spectrometry (LC-MS/MS) at multiple points over a 4 h incubation period. Results are depicted as mean ± standard deviation (SD) of three technical replicates

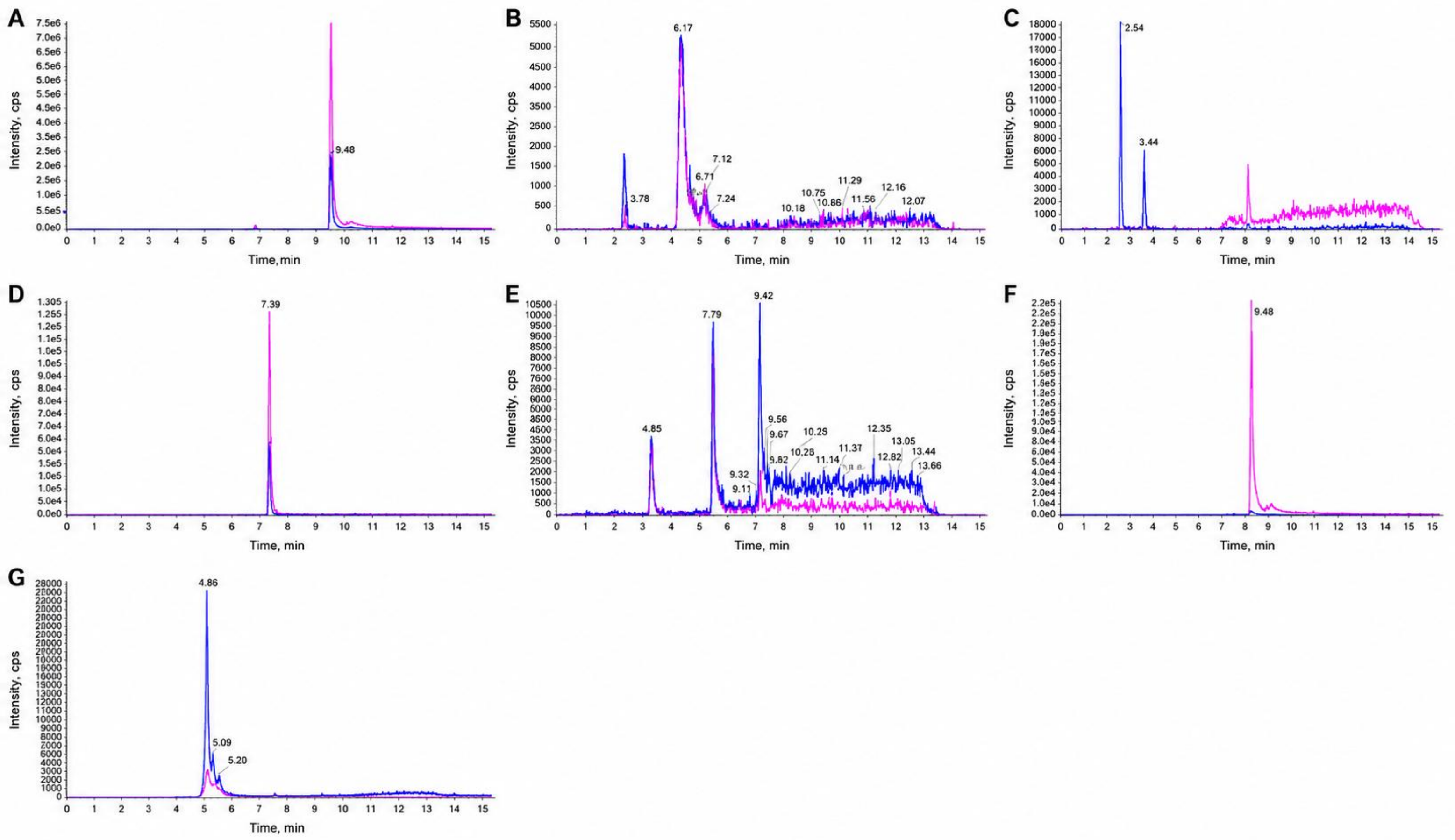


**Supplemental Figure 5.** Representative chromatograms of AME (A), AME glucuronide metabolites (B), O-demethylated AME metabolites (C), AME sulfate metabolites (D), AOH (E), hydroxylated AOH metabolites (F), and AOH sulfate metabolites (G) detected in rat hepatocytes following incubation with 8 µM AME. Chromatographic traces represent the scheduled multiple reaction monitoring (MRM) transition for AME and the corresponding MRM transitions for the respective metabolites. In-source fragmentation of AME-derived metabolites to AME was observed at their corresponding retention times.

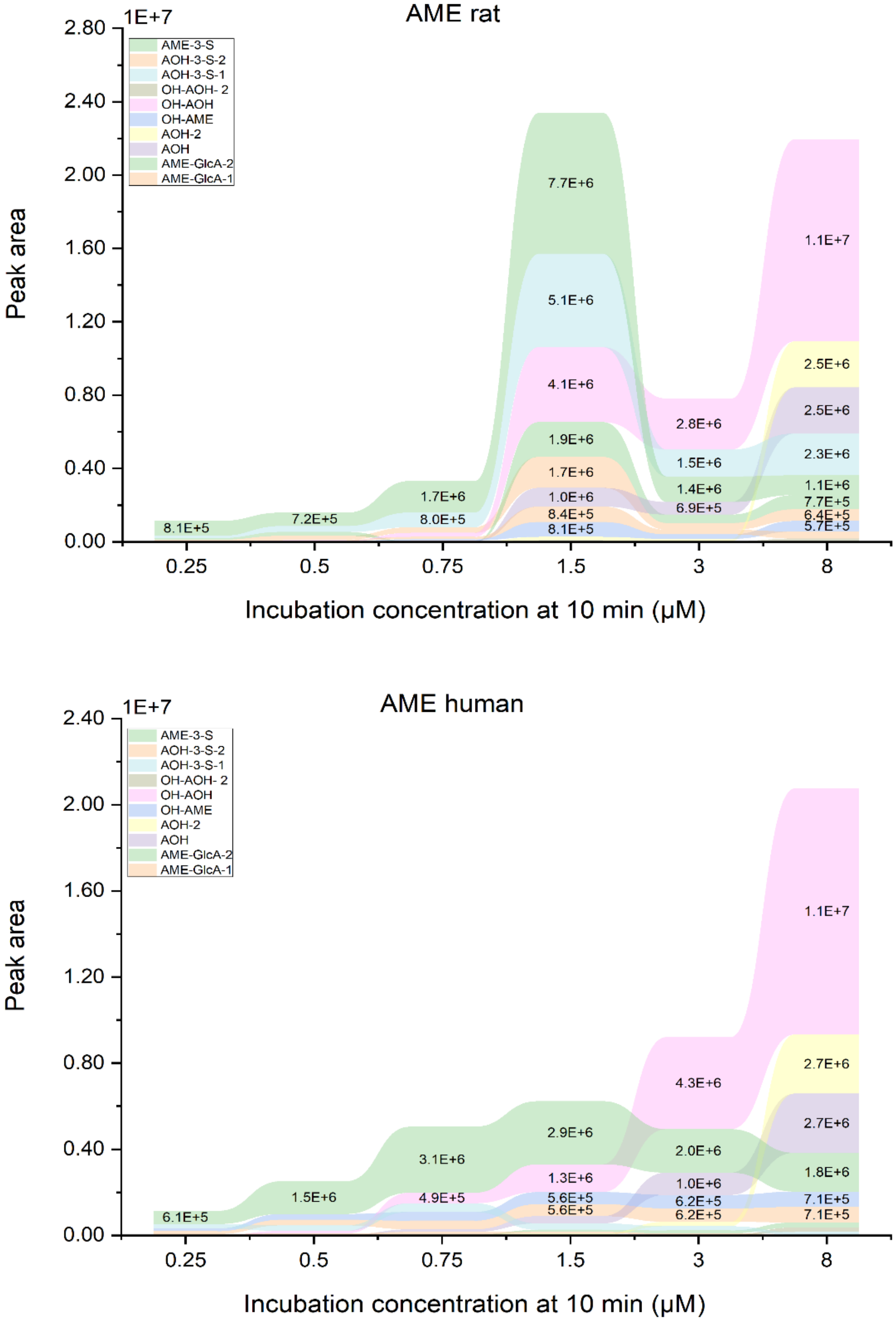


**Supplemental Figure 6**: Interspecies differences in the metabolite profile in primary rat and human hepatocytes occurring at 10 min of incubation measured via the targeted LC-MS/MS method. Each section shows the average of three independent experiments and metabolites are depicted in the order of their peak area.

**References:**

Mahmoud, M.M., Abdel-Razek, A.S., Soliman, H.S.M., Ponomareva, L.V., Thorson, J.S., Shaaban, K.A., Shaaban, M., 2022. Diverse polyketides from the marine endophytic *Alternaria* sp. LV52: Structure determination and cytotoxic activities. Biotechnol. Rep. 33, e00628. https://doi.org/10.1016/j.btre.2021.e00628
